# Phytoplankton tune local pH to actively modulate circadian swimming behavior

**DOI:** 10.1101/2023.07.24.550407

**Authors:** Arkajyoti Ghoshal, Jayabrata Dhar, Hans-Peter Grossart, Anupam Sengupta

## Abstract

Diel vertical migration (DVM), the diurnal exodus of motile phytoplankton between the light- and nutrient-rich aquatic regions, is governed by endogenous biological clocks. Many species exhibit irregular DVM patterns wherein out-of-phase gravitactic swimming–relative to that expected due to the endogenous rhythm–is observed. How cells achieve and control this irregular swimming behavior, and its impact on biological fitness remain poorly understood. Combining local environmental monitoring with behavioral and physiological analyses of motile bloom-forming *Heterosigma akashiwo* cells, we report that phytoplankton species modulate their DVM pattern by progressively tuning local pH, yielding physiologically equivalent yet behaviorally distinct gravitactic sub-populations which remain separated vertically within a visibly homogeneous cell distribution. Individual and population-scale tracking of the isolated *top* and *bottom* sub-populations revealed similar gravitactic (swimming speed and stability) and physiological traits (growth rate and maximum photosynthetic yield), suggesting that the sub-populations emerge due to mutual co-existence. Exposing the top (bottom) sub-population to the spent media of the bottom (top) counterpart recreates the emergent vertical distribution, while no such phenomenon was observed when the sub-populations were exposed to their own spent media. A model of swimming mechanics based on the quantitative analysis of cell morphologies confirms that the emergent sub-populations represent distinct swimming stabilities, resulting from morphological transformations after the cells are exposed to the spent media. Together with the corresponding night-time dataset, we present an integrated picture of the circadian swimming, wherein active chemo-regulation of the local environment underpins motility variations for potential ecological advantages via intraspecific division of labor over the day-night cycle. This chemo-regulated migratory trait offers mechanistic insights into the irregular diel migration, relevant particularly for modelling phytoplankton transport, fitness and adaptation as globally ocean waters see a persistent drop in the mean pH.

**One sentence summary:** Active regulation of local pH diversifies the diel vertical migration of motile phytoplankton.

## Introduction

Driven by the diurnal variation of light cues, phytoplankton show a periodic migratory behavior whereby they swim up to the photon-rich upper layers of the ocean during the day and swim down to the nutrient-rich deeper waters at night **[1-3]**. The ability to execute this diel vertical migration (DVM) over a 24 h day-night cycle is a fundamental trait of motile photosynthetic phytoplankton species. In combination with changes in the nutrient and the hydrodynamic conditions, DVM is critical for the fitness, succession and adaptation of phytoplankton to fluctuations in their environment, tailoring ecological niches suitable for microorganisms with diverse phenotypic spectrum **[4-7]**. Light and its gradient are key determinants of phytoplankton behavior and physiology, they impact both the swimming direction (against or toward the gravity vector) and the stability of the swimming motion under dynamic environmental conditions **[8, 9]**. Non-uniform distribution of light can further impact phytoplankton swimming, even drive populations off their regular DVM patterns **[6]**. An interplay of the intrinsic circadian rhythm and local environmental factors can engender asynchronous vertical migration and distribution of motile phytoplankton **[10-12]**. Recent studies have shown that phytoplankton actively employ exquisite mechanisms to adapt their gravitactic swimming across diverse timescales, for instance as a response to turbulent cues **[7, 13]** or under long-term nutrient limitation **[14]**.

Vertical migration and photophysiology of phytoplankton are intrinsically coupled, mediated by the internal levels of biochemicals including nitrate, carbon and other secondary metabolites, which in turn affect the propensity of vertical swimming **[10, 12]**. Depending on the species at hand, vertical migration is impacted by factors including photoacclimation **[15]**, swimming speed **[16]**, and the assimilation and regulation of nutrients by the phytoplankton species **[17]**. This characteristic diel migratory behavior of phytoplankton, known to originate solely due to changes in the light intensity, has also been reported in the deep seas where light is absent **[18]**. Thus, phytoplankton DVM is regulated by a combination of external light cues and endogenous circadian clock, as demonstrated in the case of the model unicellular species, *Chlamydomonas reinhardtii* **[19]**. Endogenous periodicity associated with phototaxis and nitrogen metabolism **[19, 20]** were reported, indicating the presence of circadian genes, both at the level of mRNA **[21, 22]** as well as DNA **[23]**. These studies show that periodic response can occur due to a variety of factors, even in the absence of a light cue. Yamazaki and Kamykowski’s model **[12, 24]**, based on cross-talks between the internal (chemical) state and changes in the light environment, suggest emergence of irregular migratory patterns and their potential adaptive advantages, particularly when nutrient availability (nitrate) changes. This model successfully explains DVM in the framework of decision-making, providing insights into the irregularities in phytoplankton migration, observed both in the lab and in field **[2, 25, 26]**.

By controlling several internal factors, phytoplankton may ‘decide’ how to respond to a particular cue **[11, 12]**. Ideally such active decision-making could assist phytoplankton to either escape a hostile environment **[7]**, or utilize the resources available optimally **[14, 27]**, thereby ensuring adequate fitness. Studies investigating the nature, cause and the effects of such active decision making–leading to out-of-phase DVM patterns–are missing this far. Lab-based experiments **[3, 28, 29]**, and proposed models have attempted to establish the decision-based vertical migration of phytoplankton **[12]**, however the mechanistic underpinnings remain unknown. While irregular migratory behavior may potentially compromise cellular photosynthesis, it may still be physiologically beneficial over different timescales and under different environmental settings **[12, 24]**. Currently, the cost-to-benefit tradeoffs of irregular DVM patterns remain to be assessed, thus leaving open a fundamental gap in our understanding of this long-observed trait, and its potential ecological role as an alternative to the circadian DVM patterns.

Here, using *Heterosigma akashiwo* (HA), a harmful algal bloom forming raphidophyte **[30-32]** as a model organism, we report that phytoplankton species modulate their DVM pattern by progressively tuning the local pH, generating behaviorally distinct gravitactic sub-populations which localize at the upper (*top* sub-population) and lower (*bottom* sub-population) regions of the cell culture. The sub-populations remain vertically separated when they are co-existing, however, in isolation, the sub-populations possess similar behavioral (swimming speed and stability) and physiological traits (growth rate and maximum photosynthetic yield), suggesting that the sub-populations emerge due to mutual co-existence. We confirm this by exposing each of the sub-populations to the spent media of their counterpart, recreating the observed vertical distribution; and through single-cell imaging, we identify morphological transformations of the cells as the cause for the distinct vertical distributions. A model of swimming mechanics reveals distinct swimming stabilities which emerge due to the morphological transformations after the cells are exposed to the spent media. The night-time data set confirmed that the sub-populations switch their behavioral and physiological traits (relative to the day-time values), indicating that phytoplankton actively chemo-regulate their local environment to promote intraspecific division of labor over a 24 h time window.

## RESULTS

### Sub-populations with similar phenotypes yet distinct swimming properties emerge within phytoplankton cell culture

We grew *Heterosigma akashiwo* CCMP452 (henceforth referred to as HA452) in culture tubes under 14:10 day: night cycle and uniform light conditions (Materials and Methods, Fig. 1A inset, top-left). The cell cultures used here originate from a monoclonal culture, which are propagated for multiple generations as continuous cultures. For the propagation, cells from the top 0.5 mm of the mother culture are collected and put in a fresh nutrient-rich medium to grow (Materials and Methods). To check if the cells grow optimally, we extracted the growth curve by measuring the cell numbers using microscopy and image processing, and compared it with previously reported growth curves of HA452 **[7, 14]**. The computed cell concentration is plotted as a function of time and shown in Fig. 1A, B. From this growth curve, the doubling time is calculated (Fig. S1), which turns out to be 26 ± 2 h, indicating that the cells have optimum growing conditions. The bottom-left inset of Fig. 1A shows a typical culture tube at 96 hours of growth, imaged during daytime (11 h). We observe that, despite optimal growth conditions and absence of any external stimulus, cells distribute all along the vertical height of the culture tube rather than confining only at the top, as expected from their diel migratory behavior **[3, 32]**. The light field in all our experiments are maintained uniform in all our experiments. While irregular patterns of DVM are known, they are usually in response to fluctuations in the environment, mostly in terms of nutrient availability. Naturally, we wanted to investigate if the distribution of nutrients along the culture tube is uniform or not. We collected cell suspensions from the top and bottom of the culture tubes at 96 h, filtered out the cells and computed the nitrate and phosphate concentrations (the two most crucial nutrients reported in irregular DVM) of the filtrate (Fig. 1C, D). The detailed steps of generating cell filtrate are shown in Fig. S2 and the same protocol is used for all experiments mentioned. We noted that there was no major difference between the concentrations of either nutrient across top and bottom samples (Fig. 1C, D). In case of a pure medium, that formed our control (Fig. 1C, D, left panel), there was also no difference between top and bottom samples, and showed comparable growth rates (Fig. 1C) suggesting homogeneous distribution. Even though there was no notable difference across samples from growing culture, it is to be noted that the overall nutrient levels went down (as compared to pure medium). This is due to active consumption by growing and dividing cells and the pattern of depletion was similar to other *H. akashiwo* strains **[14]**. Also, this points to the fact that even with cells present, distribution of nutrients remains homogeneous. We hypothesized that a difference in swimming behavior can lead to the emergence of spatial segregation even under DVM, for example directionally stable cells or cells with higher swimming speeds will be able to swim up/down and occupy these regions before the others. Towards this, we collected cells from top and bottom and investigated key static and dynamic motility parameters. First, we wanted to see the steady-state vertical distribution of top vs bottom cell samples (static parameter). We put the cell suspensions in a milli-fluidic chamber, referred to as ‘small chamber’ henceforth, (10mm x 3.3 mm x 2mm) and let them acclimatize for 20 min., and captured the distribution from short real time videos (Fig. 1E, F). The image analysis pipeline to obtain these parameters from raw data is shown in the supplementary Fig. S2. The individual vertical distribution profile for each replicate of both sub-populations is shown in Fig. S4 A, B. Ideally, cells collected from the top of the culture tube should mostly be swimming to the top of the chamber and vice versa. We analyzed the frames to obtain individual cell centroid coordinates and arranged them to numerically represent the vertical distribution seen in the frames (Fig. 1G). We then quantified the distribution using a metric called *upward bias index*, as described by Sengupta *et. al*. **[7]** (Fig. 1G, inset), *r* = (*ƒ ↑ −ƒ ↓*)/(*ƒ ↑ +ƒ ↓*) where *ƒ ↑* and *ƒ ↓* are the cell concentration in the top and bottom of an observation window. In our case, we defined them to be the top and the bottom 1/5^th^ of the chamber height (660 μm), shown by orange and lavender lines (Fig. 1E-G). For top samples, the mean *r* (± standard deviation) value across all replicates is 0.33 ± 0.16 and for bottom ones, it is 0.33 ± 0.13. Both the distribution pattern and the *r* values suggest that the behavior of top and bottom cells in isolation is identical in terms of vertical distribution i.e., under the same relaxation time, i) majority of cells in both samples swim up (*r* > 0), ii) bottom cells can swim up just as well as top cells (comparable *r* values) and iii) not all cells collected from the top of the culture tube can make it to the top of the small chamber (*r* < 1). One possibility is that the relaxation time is large enough for cells in both the samples to swim to the top even though they have naturally different swimming speeds. To test this, we quantified the speeds by flipping the small chamber by 180 degrees after 20 min. of relaxation time. As we saw from the vertical distribution, most of the cells had swam to the top of the small chamber and formed their steady-state distributions (Fig. 1G). Cells reaching the top must be strongly negatively gravitactic and after a 180 degree flip, they are at the bottom of the chamber and will naturally try to swim to the top, giving us quantitative insights into the swimming behavior (revealed by the trajectories). Note that this is true for all other cells of a sample as well, not just negatively gravitactic ones. Less negatively gravitactic cells will be present towards the bottom of the chamber and vice versa and contribute to the complete steady state distribution of the sample. We analyzed the trajectories (Materials and Methods) of both samples to obtain the mean (± standard deviation) absolute speeds for the top (85.6 ± 21.5 μm/s) and bottom (86.7 ± 23 μm/s) as shown in Fig. 1H. The samples show no significant difference, and speed can be ruled out as a major factor to trigger opposite gravitactic behavior. We also computed the *x*- and *y*-velocity components for both the samples (Fig.S5). There is no notable difference, indicating that the swimming pattern of both samples are similar. The preferred direction of swimming of a cell depends on the stability of the cell towards or away from gravity vector (biophysically speaking) and difference in stability can govern the physical location of cells along the vertical and hence the final distribution. To test this, we computed the orientational stability of top and bottom samples, with respect to the negative gravity vector (Fig. 1I). The individual plots for each replicate of both sub-populations is shown in Fig. S6 A, B. Stability is quantified in terms of reorientation time, τ_*r*_= 1/2A (A being the stability parameter obtained by plotting the instantaneous rotation rate, ω, as a function of the instantaneous angular position, *θ*. which, for the top sub-population is 12.2 ± 4.9 s and 11.48 ± 5.3 s for the bottom counterpart. These findings confirm that under uniform growth conditions, cells show identical behavior in isolation, however this does not explain the collective behavior within a cell culture. One factor could be that some cells are better at sustained up-swimming (over longer vertical distances) and others not, thus some are programmed to stay up and vice versa. We test this hypothesis next by studying the swimming behavior within a taller vertical chamber, described in the following section.

**Fig. 1.**
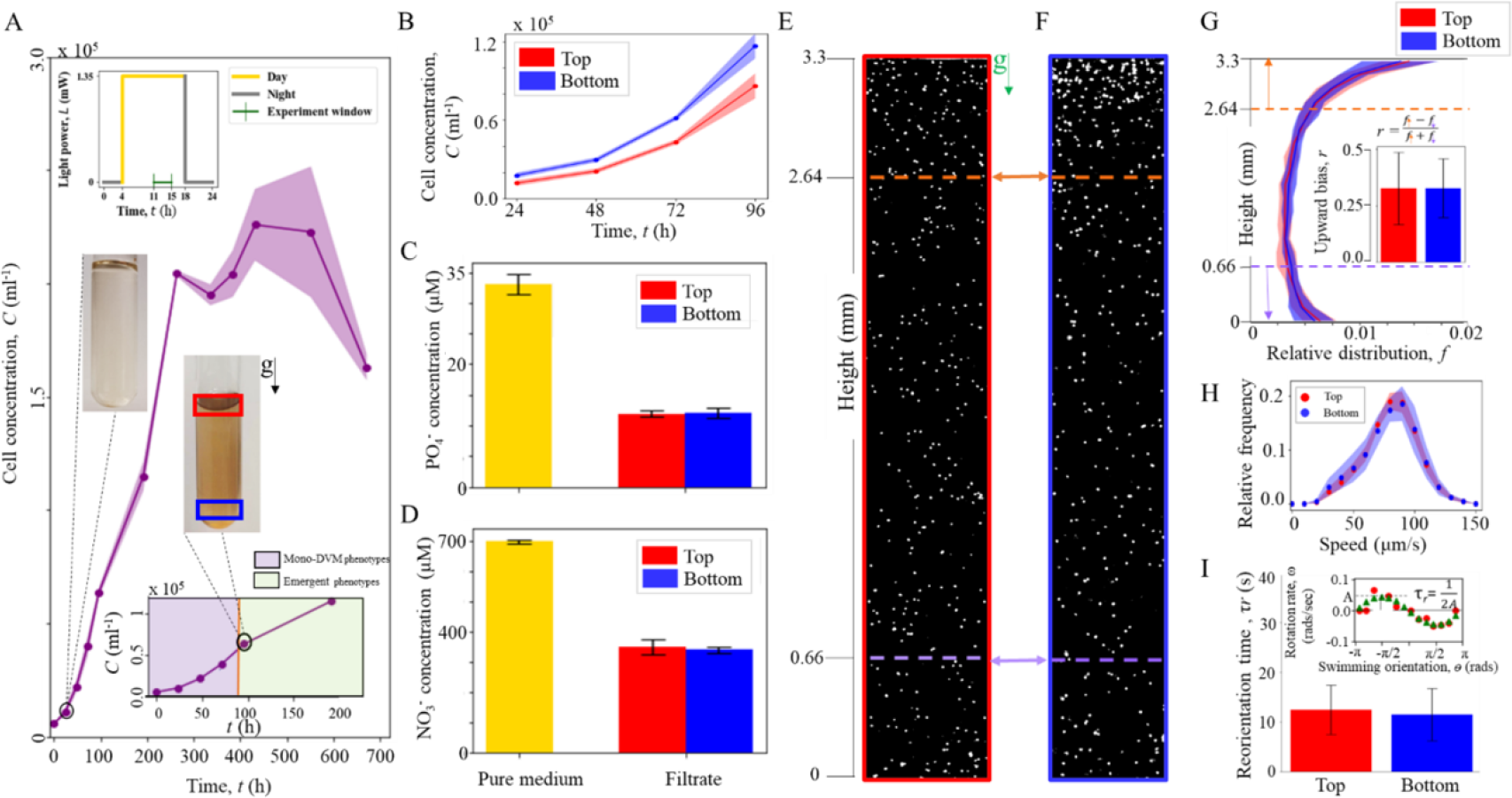
Vertically segregated sub-populations, with similar behavioral traits, emerge at the top and bottom regions of a homogeneous cell culture. **(A)** Growth curve of *Heterosigma akashiwo* (HA) 452 culture (number of replicates, *n* = 4), grown under a 14 h (light) – 10 h (dark) cycle (inset, top-left) at 22°C within a vertical culture column is ∼70mm. The inset (bottom-right) marks the timepoint at which the vertically segregated sub-populations emerge, visualized by the uniform hue of the homogeneous cell culture (upper inset panel of culture at 96 h after inoculation), compared to the vertical gradient of hue observed for the negatively gravitactic cell population (before 96 h). The darker hue on the upper part of the culture and lighter below indicate the presence of higher number of cells near the top. **(B)** Growth curves of the top (red) and bottom (blue) sub-populations, after they have been isolated from the mother culture, show similar trends relative to each other, and with that of the original culture (shown in panel **(A)**). The error (s.d.) is shown by the shaded portion of the plot. **(C, D)** Spectrophotometric analysis of the pure and spent media (96 h after inoculation) reveals comparable nitrate (C) and phosphate (D) concentrations present in the top (red) and bottom (blue) sub-populations. Yellow bar indicates nutrient levels of a pure medium. The plots show mean (± s.d.) for 2 biological replicates per sample. **(E)** Stationary distribution of the HA452 cells isolated from the top and bottom **(F)** sub-populations within a vertical chamber (height ∼3.3 mm, bright spots indicate cells, shown after subtraction of the background intensity) after 20 minutes of equilibration time. The gravity vector is indicated by **g. (G)** Stationary vertical distribution obtained from all frames of video micrography, shown over the chamber height for top (*n* = 6, average cell count per replicate, *N* = 1356 ± 283) and bottom (*n* = 6, *N* = 1757 ± 734) sub-populations. Inset shows comparable upward bias index, *r* = (*f↑ − f↓*)/(*f↑* + *f↓*) for each case, where *f↑* and *f↓* are the concentrations of cells within the first and the last 660 μm of the chamber, indicated by the arrow heads next to the dashed lines up (*↑*, dark orange) and down (*↓*, lavender) in panels (E) and (F). **(H)** Relative distribution of swimming speeds of the sub-populations, obtained by analyzing swimming trajectories (after flipping the chamber by 180 degrees), shows similar motility (*n* = 6, *N* = 670 ± 122, speed = 85.6 ± 21.5 μm/s) and bottom (*n* = 6, *N* = 829 ± 421, speed = 86.7 ± 23 μm/s). **(I)** Swimming stability measured as the reorientation timescale, τ_*r*_, (inversely proportional to swimming stability) of the top (*n* = 12, *N* = 586 ± 133) and bottom (*n* = 12, *N* = 742 ± 255) sub-populations are shown respectively as red and blue bar plots. For each cell undergoing reorientation, trajectory is analyzed to find out instantaneous angular direction of swimming (θ) and the instantaneous rotational rate (*ω*) is plotted against *θ* to obtain a curve shown by red dots in the inset. This is then fitted to a sinusoidal curve (green dots), the peak of which is the maximum rotation rate, *A*, and 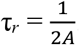. The mean reorientation timescales for the top and the bottom sub-populations are 12.8 ± 5 s and 11.5 ± 5 s respectively.

### Swimming behavior of sub-populations remains consistent over longer vertical distance

In oceans, diverse phytoplankton species can grow up to depths of tens up to hundred meters below the sea surface **[33-35]**. Swimming over large distances can be energetically costly, thus it is imperative to observe if phytoplankton alter their swimming behavior over longer vertical distances. Specifically, it would be relevant for us to check if, and how the swimming speeds of cells from the top and bottom sub-populations emerge over time, as they swim over larger vertical distances (over shorter distances, no difference was observed as reported in Fig. 1). We use a ‘tall chamber’ with a vertical height of 60 mm (Fig. 2A, see Materials and Methods section), comparable to the entire vertical depth that the cells experience within the culture tubes (the height of the culture is ∼70 mm), and measure the vertical distribution of sub-populations over time. Cells collected from the top and the bottom 0.5 mm of a culture tube were introduced to the tall chamber and allowed ∼20 min. to acclimatize within it, forming a steady distribution. Considering an average swimming speed of 85 μm/s for either sub-population, a cell with an average swimming speed would ideally reach from one end to the other (distance of 60 mm) in roughly 12 min. So, the waiting time of 20 min. should be sufficient for the strongly negative gravitactic cells to reach the top of the chamber.

**Fig. 2.**
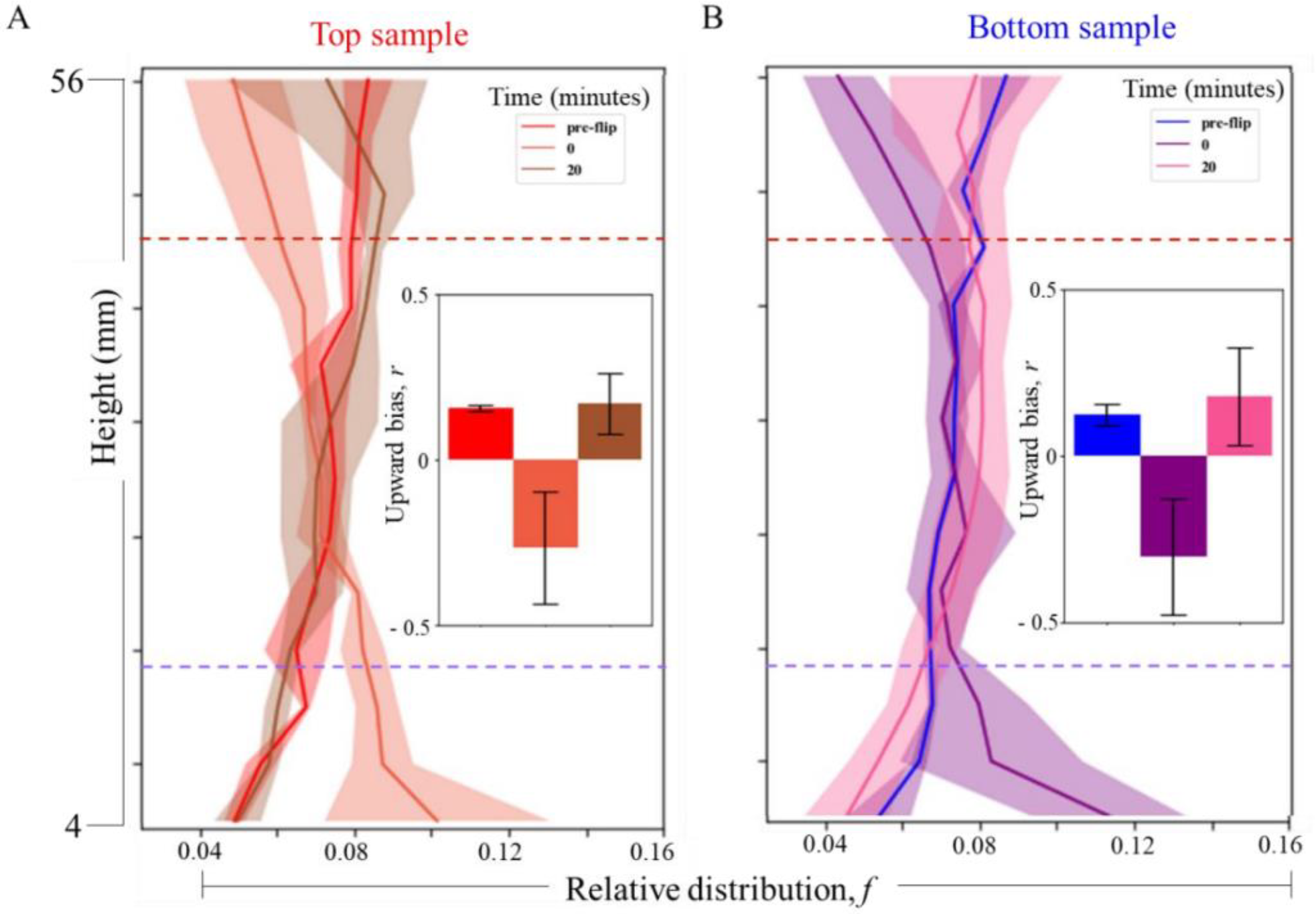
Distinct gravitactic sub-populations emerge also in longer vertical columns. **(A)** Mean ± s.d. of the vertical distribution of the top subpopulation within a 60 mm tall chamber, imaged 20 min after the chamber is filled (pre flip, *n* = 2, *N*= 35503 ± 383, red line). The chamber is then rotated by 180 degrees along the vertical plane, the distribution is recorded immediately thereafter (post flip, 0 minutes, *n* = 3, *N* = 27666 ± 3091, orange line); and after 20 minutes (post flip, 20 minutes, *n* = 3, *N* = 32563 ± 1070, brown line). The inset compares the upward bias, *r*, from the corresponding distributions (mean ± s.d.). At *t* = 20 min, pre flip, *r* = 0.157 ± 0.01; at *t* = 0 min post flip, *r* = -0.265 ± 0.17; and 20 min after flip, *r* = 0.17 ± 0.1. **(B)** Mean ± s.d. of the vertical distribution of the bottom subpopulation within the 60 mm tall chamber, imaged 20 min after the chamber is filled (pre flip, *n* = 2, *N* = 35552 ± 922, blue line). The chamber is then rotated by 180 degrees along the vertical plane, the distribution is recorded immediately thereafter (post flip, 0 minutes, *n* = 4, *N* = 31234 ± 5560, purple line); and after 20 minutes (post flip, *n* = 3, *N* = 31483 ± 5785, magenta line). The inset compares the upward bias, *r*, obtained from the corresponding distributions (mean ± s.d.). At *t* = 20 min, pre flip, *r* = 0.126 ± 0.03; at *t* = 0 min post flip, *r* = -0.3 ± 0.18; and 20 min after flip, *r* = 0.18 ± 0.15.

Thereafter, the chamber was flipped instantaneously, allowing cells to reorient and swim up against the gravity. After the flip, the entire chamber was imaged immediately (time, *t* = 0 min.) with a custom-built series-connected translation stage allowing long distance movements (Materials and Methods). Then, another 20 min. were allowed before imaging the chamber again (*t* = 20 min.). The imaging was done without the first and last 4 mm as these regions were too close to the inlet ports, where cell counting was technically challenging.

Cell distribution was considered within the remaining 52 mm (dotted line, Fig. 2A). Figure 2A, B show the distributions at 0 and 20 min. post-flip, for the top and the bottom sub-populations respectively. Once again, we found that the vertical distributions of the sub-populations in the tall chamber appear statistically comparable, indicating that the bottom sub-population, in isolation, have similar swimming characteristics as their top counterparts, even over long distances.

The similarity of swimming traits is also confirmed by the upward bias values as plotted in the insets of Fig. 2A, B (*r* = 0.17 ± 0.1 for top, and 0.18 ± 0.15 for the bottom sub-populations). Taken together, the results indicate that, when isolated, cells from the bottom sub-population can effectively swim up like the top ones, even over longer distances. Since we do not see this occur in the culture tubes, we infer that either (i) the fluxes of the top and bottom sub-populations maintain a dynamic equilibrium, shuttling continuously across the entire culture tube; or (ii) when the top and the bottom sub-populations co-exist, difference in the swimming properties emerge, relative to those in isolation. Continuous swimming to maintain a dynamic equilibrium within the cell culture is an energetically expensive option, particularly under the uniform light and nutrient conditions present in our experiments. Thus, we focus on the second possibility that distinct swimming properties emerge when the sub-populations co-exist. In the following sections, we delineate how the top and bottom swimmers emerge, and provide a mechanistic understanding of this behavior, commensurate with other physiological markers including photophysiology over both day and night-times.

### Co-existence alters swimming behavior of sub-populations via morphological changes

We observe the vertical distribution of each sub-population over a long time (120 min.), and compare this distribution of a mixed population (50:50) over the same time span. A timescale of 120 min. should be sufficient to observe any behavioral change that can alter changes in the vertical distribution (reorientation timescale ∼10 s, with swimming speed of average 85 μm/s, cells take ∼40 s to cover the vertical span of the short chamber (3.3 mm), adaptive response to external stimuli ∼10 s of seconds **[7, 13, 36]**. Fig. 3 A, C and E plot the vertical distribution over time for the isolated (top and bottom) sub-populations and their 50:50 mixture. Our results confirm that the top (Fig. 1A) and bottom (Fig. 1B) sub-populations show similar vertical distributions over time, for the entire observation window (Fig. 3 B, D). Starting with a uniform distribution at *t* = 0 min., cells occupy the upper parts of the chambers at *t* = 20 min., confirming the behavior reported in Figs. 1 and 2. Over longer durations, cells start to swim downward, reaching *r* ∼0 (or < 0, upward bias values) at *t* =120 min. While the trends we observe for the top and bottom sub-populations are similar, the rate of change of the upward bias, *r*, are different, with the bottom sub-populations showing a relatively lower rate of change compared to the top sub-population. When both sub-populations are mixed, mimicking the co-existence case, the vertical distribution pattern emerges slowly, with *r* remaining positive during the entire duration of observation (Fig. 3F). This indicates that cells from a sub-population, in presence of its complementary counterpart, tend to maintain the vertical segregation (indicated by the positive *r* values). In contrast, the isolated sub-populations switched their equilibrium swimming direction downward (Fig. 3B, D). Overall, we conclude that, in isolation, cells from both the sub-populations show similar swimming behavior, equilibrating to low *r* values over long time; whereas under co-existence, the mixed population maintains a significant positive *r* value over long times.

**Fig. 3.**
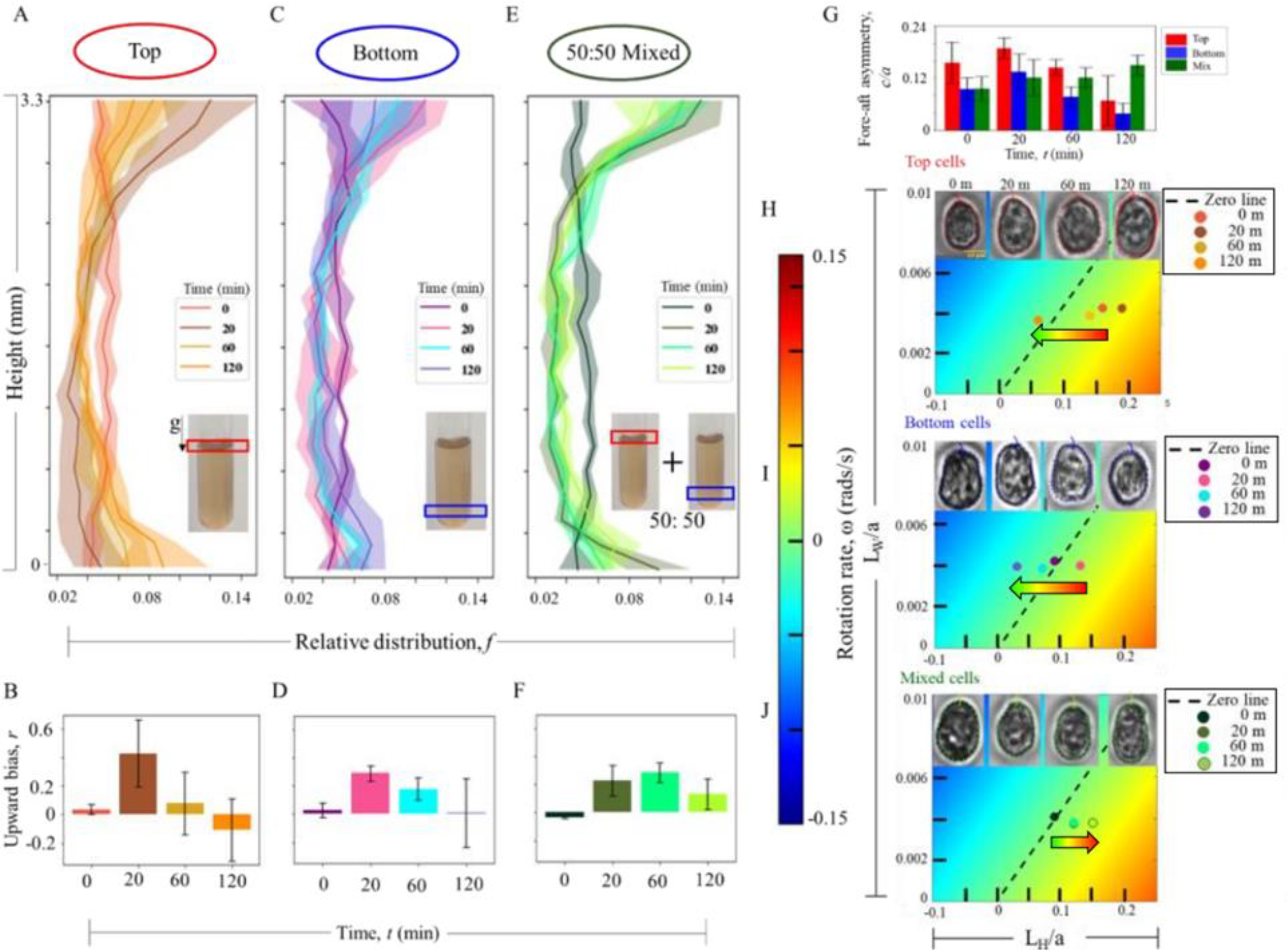
Coexistence of sub-populations triggers spontaneous alteration of cell morphology and swimming stability. Vertical distribution over time spanning 0 min., 20 min., 60 min., and 120 min., for the **(A)** top; *n* = 4, *N* = 1146 ± 205, **(C)** bottom; *n* = 4, *N* = 947 ± 306, and **(E)** 50:50 mixed; *n* = 4, *N* = 1079 ± 246 populations of cells. *t* = 0 min corresponds to start of the imaging immediately after the chamber was filled. The corresponding upward bias values (*r*) over time is shown in panels **(B), (D), (F). (G)** Morphological analysis of single cells reveal change in the fore-aft shape asymmetry over time (indicated by the value of *c/a*) for the three cases; bar plots show mean ± s.d. **(H)**. Single cells were imaged in each case (*N* = 10), from which the cell contours were extracted and fitted to a mathematical curve (Materials and Methods) to obtain cell size parameter, *b/a* and the fore-aft asymmetry parameter, *c/a*, where *a, b* and *c* are the major, minor axes and degree of fore-aft asymmetry respectively. A cell mechanical model captures the emerging variation of swimming stability (orientational stability) over time, as *L*_H_/*a* and and *L*_W_/*a* values change. *L*_H_ is the distance between the centre of buoyancy and the centre of hydrodynamic stress, while *L*_W_ is that between the centre of buoyancy and the centre of mass (see Materials and Methods). Varying one with respect to the other alters the orientational stability of the swimming cell, which is captured by the rotation rate, *ω*: *ω* > 0 (< 0) indicates cells which are stable against (toward) the gravity direction; *ω* = 0 indicates neutrally stable cells with no preferred direction of orientation (dotted black line). When isolated, the orientational stability of the top and the bottom sub-populations reduces over time (arrow indicates leftward as time increases, top and middle panels), whereas for the mixed sample, orientational stability increases over time, while remaining positive throughout (bottom panel). The insets at the top of each panel show representative single cell images for each case, obtained using phase contrast microscopy.

To find the mechanistic underpinnings of the difference in the swimming trends, we quantified the morphology of the cells from each of the experimental samples. Morphology is a key parameter that determine the swimming stability and any difference therein, that could alter the swimming direction. At t = 0 min. (reflecting the native morphological state of the cells, similar to those in the culture tube), cells from top sub-population show a high degree of fore-aft asymmetry (*c/a*) compared to the bottom or mixed cases (Fig. 3G, H). Higher *c/a* indicate high degree of fore-aft asymmetry, resulting in strongly negative gravitactic swimming properties, and vice versa. As time elapsed, the *c/a* value increased for both the top and bottom sub-populations (Fig. 3H, top and middle panels), reaching 0.19 ± 0.3 for the top cells, and 0.14 ± 0.5 for the bottom cells. This indicates that cells–over short time scales following the transfer to the millifluidic device–increased negative gravitactic ability, a signature of the reduction of the physiological stress **[7, 13]**. At 60 min., both the bottom and the top cells start to get symmetric in morphology (*c/a* lowers to 0.14 ± 0.02 and 0.08 ± 0.03 respectively), with the trend continuing during the entire course of our experiments (120 min.). At 120 min, cells from both sub-populations attain similar *c/a* values (0.07 ± 0.06 and 0.04 ± 0.02), confirming that, in isolation, the top and bottom sub-populations have comparable morphotypes (Fig. S7, S8). While the top and bottom sub-populations exhibit similar variation of fore-aft asymmetry with time, the top cells, relatively, are more elliptical than the bottom ones, as captured by the *b/a* values (Fig. S10). Consequently, this results in an overall decrease in the swimming stability over time, as shown in the phase plots (Fig. 3H, upper and middle panels).

In the mixed population scenario (a 50:50 mixture of top and bottom sub-populations), the initial fore-aft asymmetry (*c/a* = 0.10 ± 0.04) was lower than that in each of the individual sub-populations. The cells show a slight increase in the *c/a* value (0.12 ± 0.04 at *t* = 20 min.), reaching up to 0.15 ± 0.04 by 120 min. With respect to the individual sub-populations, the observed trend – an overall increase of the fore-aft asymmetry – is opposite, as also corroborated by the vertical distribution, and the swimming stability captured by our computational fluid dynamical model (Fig. 3H, lower panel). Statistically, the overall change in the population fore-aft asymmetry could arise due to the increase of *c/a* values in some cells, and lowering in others, effectively retaining a uniform value (Fig. S9). Furthermore, this indicates an emergent stability, supported by the data obtained at 20 min: the *c/a* of mixed population is lower than that of the average value of the top and bottom sub-populations, thus showing that the stability of the mixed population is not simply a superposition of the swimming stabilities of the individual components. From this we can conclude that under co-existence, the mixed population shows a distinct swimming stability and vertical distribution, otherwise not observed under isolated conditions.

### Phytoplankton chemo-regulate local pH to diversify the circadian swimming behavior

The distinct alteration of the cell morphology in the 50:50 mixture, relative to the isolated sub-populations, indicates that cells in the mixture experience physiological stress that trigger morphological changes, ultimately impacting the swimming behavior **[7, 13]**. We identify the local pH, and the relative difference between the pH of the spent media associated with the top and the bottom sub-populations, as the driver of the reported observations (Fig. 4 A). As the population grows, the pH of the cell culture increased from 8.72 ± 0.03 (top, 8.82 ± 0.04 for bottom) ∼48 h after inoculation, to 9.01 ± 0.02 (9.09 ± 0.06 for bottom) at ∼96 h.

**Fig. 4.**
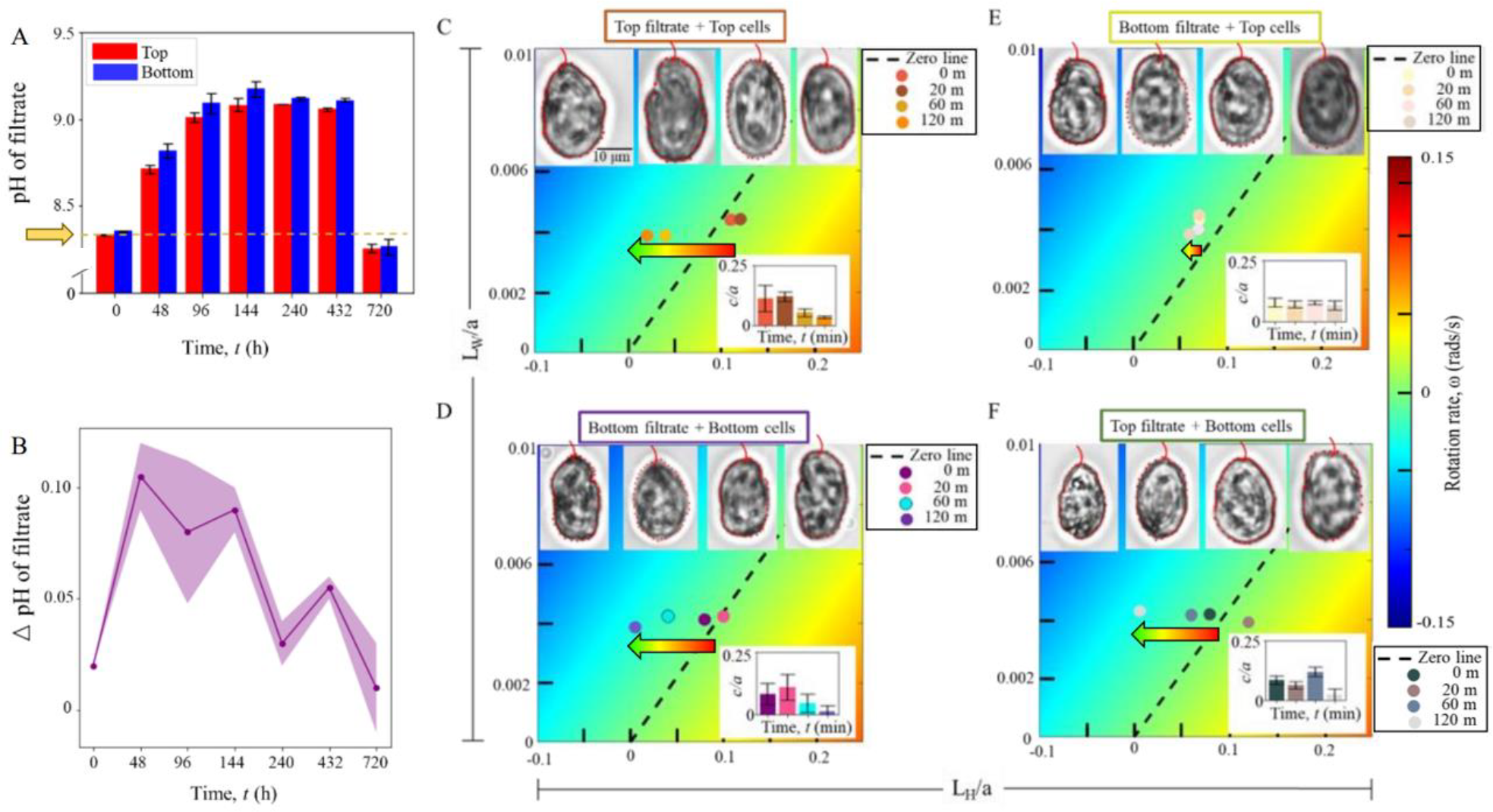
Variation of the local pH tunes gravitatic swimming. **(A)** Variation of the pH values (mean ± s.d.) over time, for the spent media obtained from the top and bottom sub-populations. *t* = 0 h corresponds to the measurement done immediately after inoculation. The pH of the pure medium is indicated by the arrow. Measurements are done on two biological replicates (*n* = 2), with multiple technical replicates in each case. **(B)** Difference of pH of the spent media (bottom minus top), corresponding to the above time points. (**C)-(F)** To capture the effect of the filtrate pH on cell populations: four different combinations of cells and spent media were designed. Corresponding effects of the spent media on the cell morphology were measured, as described previously (*N* = 10, representative single cell images are shown in panels (**C)-(F)** insets). **(C)** Represents top filtrate mixed with top subpopulation [filtrate, cell] = [*T, T*]. The bottom right inset shows *c/a* values, which change from 0.11 ± 0.06 at *t* = 0 min to 0.03 at *t* = 120 min (at *t* = 20 min and 60 min, the values are 0.12 ± 0.02 and 0.05 ± 0.02 respectively). **(D)** [*B, B*]: Bottom filtrate mixed with bottom subpopulation results; the *c/a* value changes from 0.08 ± 0.04 at *t* = 0 min to 0.01 ± 0.02 at *t* = 120 min (at *t* = 20 min and 60 min, the values are 0.11 ± 0.05 and 0.05 ± 0.04 respectively). **(E)** [*B, T*]: Bottom filtrate mixed with the top subpopulation; The *c/a* value remains nearly constant between 0.07 – 0.08 (*c/a =* 0.08 ± 0.04; 0.07 ± 0.05; 0.08 ± 0.04 and 0.07 ± 0.02 at *t* = 0, 20, 60 and 120 min). **(F)** [*T, B*]: Top filtrate mixed with bottom cells; *c/a* values are 0.08 ± 0.02, 0.06 ± 0.02, 0.12 ± 0.02 and 0.02 ± 0.03 at *t* = 0, 20, 60 and 120 min. The swimming stability shifts over time, from negative to positive gravitaxis, indicated by the arrow heads, for [*B, B*], [*T, T*] and [*T, B*] combinations (equivalent to the isolated top and bottom sub-populations, Fig. 3), while for the mixed case, [*B, T*], presence of the bottom filtrate enables the cells from the top subpopulation to maintain their morphology, thereby hindering any shift in the swimming stability.

Thereafter, the pH attains a steady value of ∼9.1, reaching a maximum of 9.16 (for both the top and bottom spent media) at 480 h, before finally dropping back to ∼8.3 at ∼720 h. This implies that as cells grow, they actively modulate the pH of their environment. Across all measurements, the pH associated with the bottom sub-population was found to be higher than that of the top sub-population (Fig. 4B), indicating the difference in local chemo-environment experienced by the cells. Comparable changes in local pH have been reported previously for native algal cultures **[37, 38]**, while small variations in exogenous stressor concentrations, e.g., via H_2_O_2_ or altering the carbonate chemistry (which also induce change of local pH) could be sufficient to impact the swimming characteristics of *H. akashiwo* **[13, 39]**.

To confirm the role of the local pH in driving the observed changes in cell morphology, and consequently, the swimming behavior, we exposed cells from each sub-population to their own (control) and to the complementary (test) spent media, thereby analyzing 4 different combinations: [spent media, cells] = [*T, T*], [*B, B*], [*B, T*] and [*T, B*], where *T* and *B* respectively denote Top and Bottom (Fig. 4C-F). The morphological data corresponding to the combinatorial experiments are presented in Figures S11-S15.

When cells from the top (bottom) sub-population were introduced with their own spent media, the fore-aft asymmetry, *c/a*, changed from 0.12 ± 0.06 (0.08 ± 0.05 for bottom) at *t* = 0 min. to 0.04 ± 0.01 (0.04 ± 0.03) at *t* = 120 min (inset, Fig. 4C, D), resulting in comparable shifts in the swimming stability of each of the sub-populations as shown in Fig. 4 C, D for the [*T, T*] and [*B, B*] combinations respectively (for both cases, the swimming stability reduces, as also observed in Fig. 3H for the isolated sub-populations). Over similar timescales, the [*T, B*] combination showed similar response (Fig. 4 F and inset): the *c/a* changed from 0.08 ± 0.03 at *t* = 0 min. to 0.03 ± 0.03 at *t* = 120 min; with an overall reduction of the swimming stability.

Exceptionally, the morphological alteration and the resulting changes in the swimming stability differs for the [*B, T*] combination, wherein cells from the top sub-population are introduced to the spent media from the bottom sub-population. In this case, no perceptible change in the cell asymmetry was observed over time (Fig. 4E and inset) with the *c/a* maintaining a steady mean value between 0.07 and 0.08. Alongside, the swimming stability remains positive throughout, indicating that the cells maintain an up-swimming behavior (against gravity) over the entire observation window, in contrast to the other three combinations. We attribute this distinct tendency of the top sub-population to remain further away from the bottom filtrate to the relatively higher pH of the spent media associated with the bottom sub-population; thus rendering the bottom filtrate relatively more potent in triggering the vertical separation of the sub-populations in the cell culture, than the other way round. Overall, the combinatorial experiments suggest that the top and bottom sub-populations emerge due to the differential response of the *H. akashiwo* population, at individual scale, to the local pH. Despite difference between the absolute *c/a* values of the top and bottom sub-populations, cells show similar swimming behavior and temporal trend under isolated conditions, which however was altered by the introduction of the spent media. The alteration of the morphological and motility traits indicates that in presence of the complementary spent media, cells–with otherwise similar traits–can elicit distinct responses which highlight the role of chemo-regulation of the local environment in tuning the phenotypic traits actively, ultimately resulting in emergent spatial distribution of sub-populations within a growing cell culture.

### Diel variation of the local pH, photophysiology and night-time swimming behavior

In order to obtain insights throughout a 24 h cycle, we carry out night-time pH measurements on the spent media associated with the top and the bottom sub-populations of a 96 h culture (Materials and Methods). As shown in Fig. 5A (right most bars), the night-time difference in pH, ∼0.02 (8.96 ± 0.04 vs 8.94 ± 0.03), is much lower than the day-time difference of the pH ∼0.37 (9.09 ± 0.06 vs 8.72 ± 0.03). While the pH of the top filtrate remains nearly constant, the pH of the bottom filtrate shows a considerable drop between the day and night-times. Alongside, at night-time, the bottom sub-population shows a significantly higher upward bias (Fig. 5B). The drop in the pH promotes downward swimming of the top sub-population, thereby aiding DVM following the circadian rhythm (Fig. 5B, upper inset). In contrast, the reduction of the reorientation timescale of the bottom sub-population (Fig. 5B lower inset, and Fig. S16), indicates that these cells are now primed to swim against gravity, thus confirming that the bottom sub-population possesses an out-of-phase circadian swimming behavior.

**Fig. 5.**
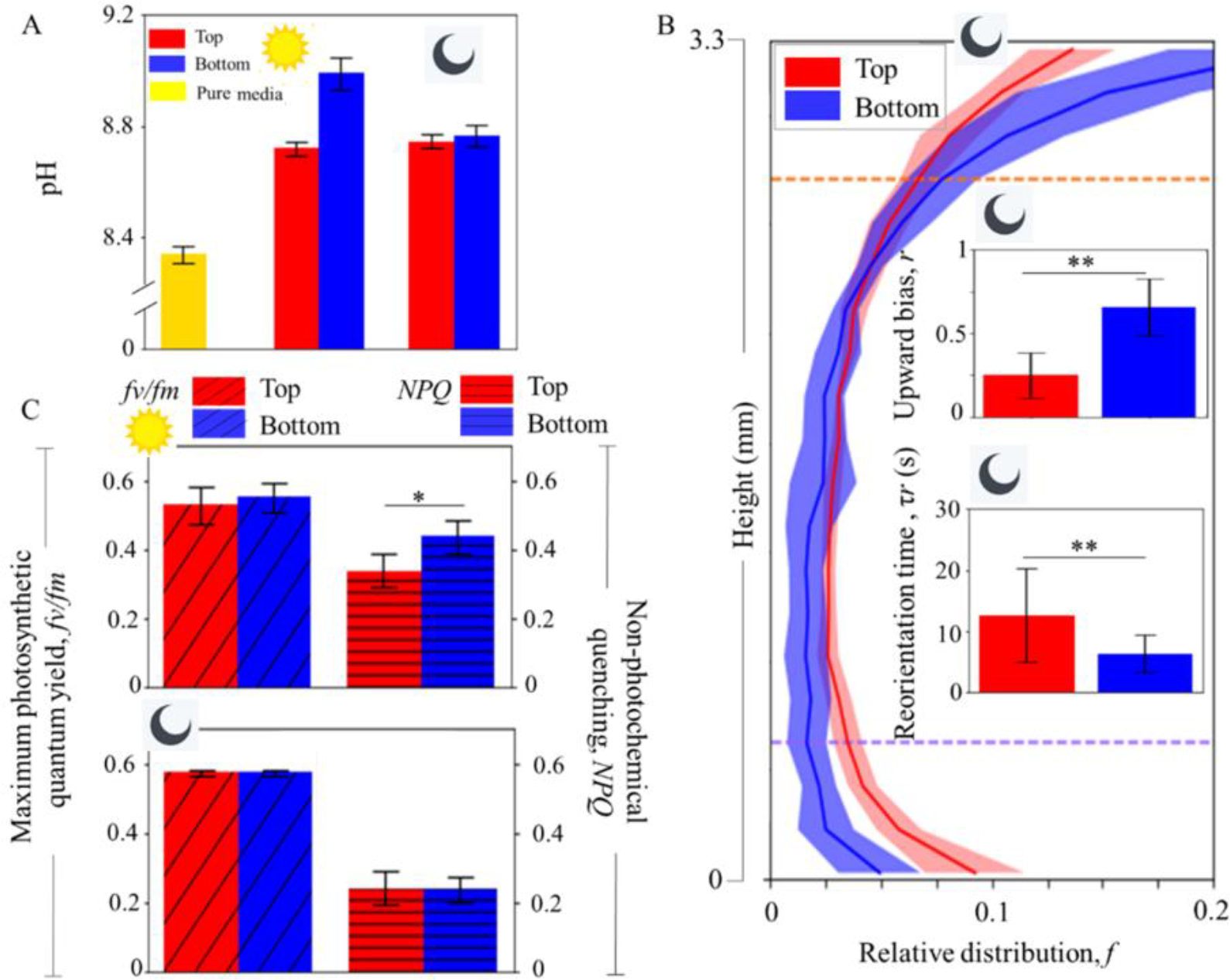
Diel variation of local pH underpins night-time swimming behaviour of gravitactic sub-populations. **(A)** Night-time pH of the spent media obtained from the top and the bottom sub-populations (20:00 h CEST, rightmost bars) 96 h after inoculation, as compared to the daytime values (11:00 h CEST, *n* = 3 in each case). The night-time difference in pH, ∼0.02 (8.96 ± 0.04 vs 8.94 ± 0.03), is much lower than the day-time difference of the pH ∼0.37 (9.09 ± 0.06 vs 8.72 ± 0.03). The pH of the bottom filtrate shows considerably higher drop (from day to night), while the pH of the top filtrate remains nearly constant. **(B)** Vertical distribution of the top (red line and shaded region show respectively the mean and s.d, *n* = 6, *N* = 1614 ± 395) and the bottom (blue line and shaded region, *n* = 6, *N*=1112 ± 418) sub-populations at night-time, around the midnight (00:00 h, indicated by a crescent moon). The upper bar plot in the inset shows corresponding upward bias index (*r*): 0.26 ± 0.14 for top and 0.66 ± 0.17 for bottom, reveal significantly higher up-swimming stability of the bottom subpopulation at night-time (p < 0.05, two asterisks, 2 tailed t-test). The lower inset plotting the swimming stability in terms of the reorientation timescale, τ_*r*_ (mean ± s.d.): 2.6 ± 6 s for the top (*n* =12, *N* = 733 ± 174) and 6.3 ± 3 s for the bottom (*n* =12, *N* = 545 ± 207) sub-populations. The timescales are significantly different, p < 0.05 (two asterisks), verified using a 2 tailed t-test. **(C)** Top panel presents the daytime (marked with a sun) maximum photosynthetic quantum yield, *fv/fm* for the top (red, *n* = 4) and the bottom (blue, *n* = 3) sub-populations after two minutes of dark adaptation. No statistically significant difference was noted (p = 0.63, 2 tailed t-test). However, the non-photochemical quenching (*NPQ)* values show significant differencet (p < 0.1, indicated by the asterisk on the bar plot on the right). In the bottom panel (night-time), the bar plots in the left show maximum photosynthetic quantum yield, *fv/fm* for top (red, *n* = 2) and bottom (blue, *n* = 2) samples after two minutes of dark adaptation. No statistically significant difference is noted (2 tailed t-test, p = 0.94), also for the corresponding *NPQ* values (p = 0.96).

Under optimal conditions, DVM patterns–mediated by the internal circadian rhythm–allows phytoplankton species access to sunlight and nutrients while avoiding detrimental factors including predators. As such, an out-of-phase pattern would emerge when the fitness benefits are limited under evolutionarily established DVM. In our case, 96 h after the inoculation, the nutrient levels are considerably reduced, and will be fully depleted within ∼200 h. Thus, it could be advantageous for a population to expand its vertical distribution to enhance the search radius for resources, as well as spread over a larger space to grow and divide without crowding. This may be achieved through turning a part of the population to swim downwards (forming the bottom sub-population), as against all the cells competing for limited resources and space by swimming up (top cells). Such an active diversification of the swimming strategy could be further rationalized if, additionally, the emergent sub-populations possessed differential photophysiology and ability to alleviate light stress (top swimmers will be, on average exposed to higher doses of light that those at the bottom). By carrying out pulse-amplitude modulated chlorophyll fluorometry (PAM) assays on each of the sub-populations, we estimate their photophysiology during the day and night-times (Fig. 5C). While there is no statistically significant difference between the maximum photosynthetic quantum yield, *fv/fm*, and the maximum electron transfer rate, *ETR*_*max*_ (Fig. S17) either during day or night, the NPQ values show a significant difference during the day-time and no difference at night time.

This indicates that the cells from the bottom sub-populations are at a higher oxidative stress level than top ones (in agreement with observations from natural systems). This may explain why the bottom cells are rounder (lower *c/a*), and can also result in production of excess extracellular ROS (e-ROS) by the bottom cells, thus rendering the bottom environment more potent as an effective info-chemical as seen in Fig. 4. When either sample was isolated and dark-adapted for 20 min., there was no notable difference in any of the photophysiological parameters, indicating that, in absence of the complementary sub-population, the photophysiological parameters are comparable as also seen for the other traits. Thus, we can conclude that even though cell samples at top and bottom have similar photosynthetic efficiency, the capacity of stress handling is different. So, it would be beneficial for some cells to stay at top and others at bottom. One key feature of vertical sub-populations is the fact that in isolation, they show effectively comparable behavior and physiology, and thus present functionally equivalent states. In natural marine settings, this may be critical for species survival especially if a portion of the population gets eliminated by deleterious stimuli like turbulence, the other sub-population can effectively function and maintain the biological fitness of the species.

## Discussion

Recent studies suggest that the production of info-chemicals (signaling molecules) by phytoplankton can regulate population-scale behavior, offering potential biological functions like avoiding predators **[40-42]**. Studies show that change in the exogenous pH can trigger change in swimming behavior and potentially alter the DVM pattern of phytoplankton **[39]**. Decrease in pH values from 8.26 to 8.13 was sufficient to induce increased downward swimming in *H. akashiwo*. In natural settings, pH variations occur due to the direct addition and removal of dissolved inorganic carbon (DIC) owing to photosynthesis and cellular respiration **[43]**. Over the course of a 24 h period, metabolic processes **[44, 45]**, in combination with hydrodynamic factors **[46, 47]**, can engender considerable changes in the local pH. Our results agree with reported observations on the impact of pH on the motility, a key difference that we note here is that the cells, as they grow, modulate their local pH to generate distinct sub-populations. The pH associated with the top and bottom sub-populations represent considerable difference in the local alkalinity experienced by the cells, resulting in distinct behavioral responses when cells are introduced to their complementary spent media (as revealed by the combinatorial experiments). This ultimately leads to the vertical separation of the sub-populations within a growing culture, with the relative difference in the local pH acting as an effective info-chemical.

An alternative relevant mode of cell-to-cell chemical communication is the quorum sensing, wherein signalling molecules called autoinducers are released when local population density reaches a critical value, particularly under confinements **[48, 49]**. So as to ascertain if the phytoplankton traits observed here emerges due to quorum sensing, we studied cells from continuous cultures at 48 h and 96 h of after inoculation (Fig. S18). At 48 h, the upward bias (mean ± s.d.) is 0.52 ± 0.27 for the top cells, and 0.48 ± 0.21 for the bottom cells. We note that in isolation, the top and the bottom cells showed similar upward bias values as for the cells which were in the 96 h culture (data shown in Fig.1). The 48 h culture had a higher upward bias than the 96 h population, indicating that younger cell cultures comprised higher proportion of up-swimming cells, in agreement with the DVM trends generally observed. As shown in Fig. 1A, B, the emergent sub-populations are observed around 96 h, when the population density is ∼3-fold relative to that at 48 h. If a quorum-like behavior triggered the emergence of sub-populations in cultures older than 48 h, diluting 96 h culture (to match the cell concentration at 48 h) should drive the upward bias values higher. However, no such enhancement of the upward bias values in the diluted 96 h cell cultures were observed (Fig.S18), confirming that the sub-populations emerge due to co-existence, and not by a high population density of the cells. The lack of a quorum-like behavior is further confirmed when we expose the sub-populations to their corresponding (or complementary) spent media as discussed in Fig. 4. This conclusively indicates that it is cue for the emergent behavior lies in the spent media, effectively acting as an info-chemical that diversifies the circadian swimming behavior.

Finally, we tested an additional mode of communication commonly found in phytoplankton communities: extracellular vesicles (EVs). EVs have been shown to mediate cell-cell communications over large distances **[50]**. While we detected low concentrations (∼10-100/ml) in our experiments, careful investigation of our samples did not however yield any observable differences in the circadian swimming behavior when we compared swimming behavior in the presence or the absence of the EVs in the local environment.

In conclusion, we have shown that a monoclonal phytoplankton population can engender behaviorally distinct phenotypes over time, due to *in situ* variation of the local pH. The emergent phenotypes are behaviorally plastic, and can play interchangeable functional roles depending on the time of the day (light vs dark periods). Although both subpopulations were found to be negatively gravitactic during daytime, one performed better than the other under co-existing conditions, thus leading to a vertical segregation of the population into strong and weak up-swimmers. Evolutionarily, cells are programmed to be up-swimmers during daytime (to get light for photosynthesis), so, those which do not show strong up-swimming behavior may compromise fitness. To circumvent this potential drop in fitness, the sub-populations develop complementary circadian swimming patterns, additionally enabling the population, as a whole, to expand their vertical niche which is advantageous for accessing resources as well as increasing the survival probability. While at a population scale, circadian swimming is harnessed to make the system robust and dynamic, at a single cell level, cells leverage environmental pH as a intra-population communication cue to trigger their vertical segregation. As the overall pH of the medium becomes basic, cells which are strong up-swimmers, are capable of reducing the pH of local environment (acidic) during day and vice versa. This in turn, reduces (pH) stress, making them more fore-aft asymmetric and better up-swimmers. Change in morphology due to stress may be associated with the production of reactive oxygen species (ROS) in *H. akashiwo* **[7, 13]**. Phytoplankton are major producers of ROS in marine environments and many species are known to produce e-ROS even under optimal growth conditions **[51]**. Production of extracellular ROS can be a possible mechanism to get rid of excess endogenous or surface ROS **[52, 53]**. While e-ROS has been attributed to functions like predator avoidance, resistance to viral infection, and toxicity in case of HABs, they have also been associated to allelopathic interactions and communication with other species **[40]**. Thus, ROS, alongside reactive nitrogen species **[42]**, can potentially act as cues for conspecific cell-cell communication and signaling. Based on their chemical forms, reactive species have the potential to alter the pH of the local media, and therefore, may be at play in mediating the emergent phytoplankton behavior we have reported here. In marine ecosystems, changes in the pH depend on the buffer capacity of the seawater, which, due to ongoing anthropogenic stresses is expected to decline in the future. This may have profound implications for food webs: not only photosynthesis of phytoplankton, but also swimming and hence photosynthetic efficiency of motile species will be directly impacted. Globally, as the ocean waters see a rapid change in the pH **[54]**, our results will provide a mechanistic basis for assessing the impacts of ocean acidification on phytoplankton migratory patterns, and shed light on the transport, fitness and adaptation of phytoplankton inhabiting oceans of the future.

## MATERIALS AND METHODS

### Cell Culture

Cells were cultured in 50-ml sterile glass tubes with 14:10 hours of light and darkness (to simulate a diel cycle) in f/2-Si (minus silica) medium at 22°C and constant light intensity (of 1.35mW, during light phase). They were propagated every 1 to 2 weeks, by inoculating 2 ml of cell suspension collected from ∼0.5 mm of the top of a parent culture (mother culture), into 25 ml of fresh medium, in a laminar flow chamber. White light (wavelength, λ= 535 nm, power =1.35 mW) illuminates the incubator between 4 hours to the end of 17 hours (yellow line) and mimics daytime. It is switched off automatically at the start of 18^th^ hour till the 4^th^ hour of the next day (gray line), to represent night. Experiments are usually performed between 11 to 15 hours, indicated by the green line region. For all experiments reported in this paper, we used cultures that were 96 hours old (early exponential phase of growth). All daytime experiments were conducted between 10 hours to 16 hours to rule out any DVM related impact. Nighttime experiments were conducted between 20 hours to 22 hours. We started 24-hour experiments at 11 hours of one day and concluded at the same time of the next day. For each study, unless mentioned otherwise, we used at least four replicates, the details of which are provided in their corresponding sections. All experiments reported here can be broadly classified into a) population scale and b) single-cell level. The experimental setup and protocol for both categories are outlined below. Unless otherwise mentioned, all experiments reported in this article uses cells that are about 96 h old. For all experiments reported in this article, cells are collected from ± 5 mm of the 2 different regions shown on the culture tube. Cells collected from the red box indicated region of the culture tube are ‘Top’ samples and the blue box corresponds to ‘Bottom’ samples.

### Quantification of growth curve

To quantify growth curve, fresh cultures were prepared from parent cultures. Cells were allowed to grow for 24 hours from whence, measurements were taken. For the 0-hour time point reading, cells were collected from the parents separately and counted. The count was then normalized assuming dilution in a 25 ml culture and from this the final count was taken. From 24 hours, cells in the cultures were counted in a 3.5 μl chamber and observed under a stereo microscope (Nikon ® SMZ1270). Readings were taken for 24, 48, 72, 96, 192, 264, 336, 384, 432, 552 and 672 hours (28 days). The counts were converted to the appropriate units and growth curve was plotted (count as a function of time). To calculate doubling time from this data, cell count was replotted in log scale against time. The time points showing linear curve (indicative of exponential growth/ doubling growth) i.e., 0 to 96 h were selected. From this, growth curve was calculated as

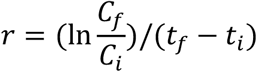

where *r* is the growth rate, *C*_f_ and *C*_i_ are cell concentrations between consecutive time points *t*_f_ and *t*_i_ (e.g. 24 and 0 h, 48 and 24 h, and so on). From this, doubling time was calculated as

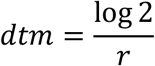

where *dtm* is the doubling time. The *dtm* of each pair of timepoints were averaged to obtain the mean (± s.d.), which is reported.

### Cell harvesting and preparing cell filtrate

For each experiment reported, unless otherwise mentioned, two types of samples were used: top samples: which are cells collected from the approximately the first 0.5 mm of the culture using 1ml pipette (+tips) and bottom samples, which are cells collected from around 0.7 mm above the base of the culture tube. To collect this, Sterican^®^ disposable long needles (120 mm x 0.8 m) were used with Injekt^®^ disposable syringes. This harvesting involves collection of cells plus their surrounding fluid medium. In experiments where just the medium was needed, cells were filtered out using 1μm (pore size) Whatman^®^ Puradisc 25 syringe filter. The detailed steps are shown visually in supplementary Fig. S2.

## Population scale experiment

### i) Quantification of vertical distribution from an equilibrium state

All vertical distributions were observed in either a ‘small’ (10 mm x 3.3 mm x 2 mm) or a ‘tall’ (60 mm x 3.3 mm x 2 mm) chamber made of polymethyl methacrylate. For small chamber experiments, a chamber was mounted on a custom cage for visualization and imaged using a lens-camera setup. The cage is equipped with *xz* translation screws, with a maximum displacement of 25 mm. The entire setup, as well the calculation methodology is detailed in **[14]**. Cells were filled into it and visualized at ∼1.7 x zoom, using a Grasshopper3 (model: GS3-U3-41C6C-C) camera with an 1″ sensor. At this zoom, the entire chamber was captured in one go. The chamber was illuminated with a red LED (630 nm wavelength). Imaging was done in the middle region (horizontal) of the chamber to stay away from the inlets and boundary walls. After filling, cells were allowed 20 minutes to reach their steady distribution which was then imaged.

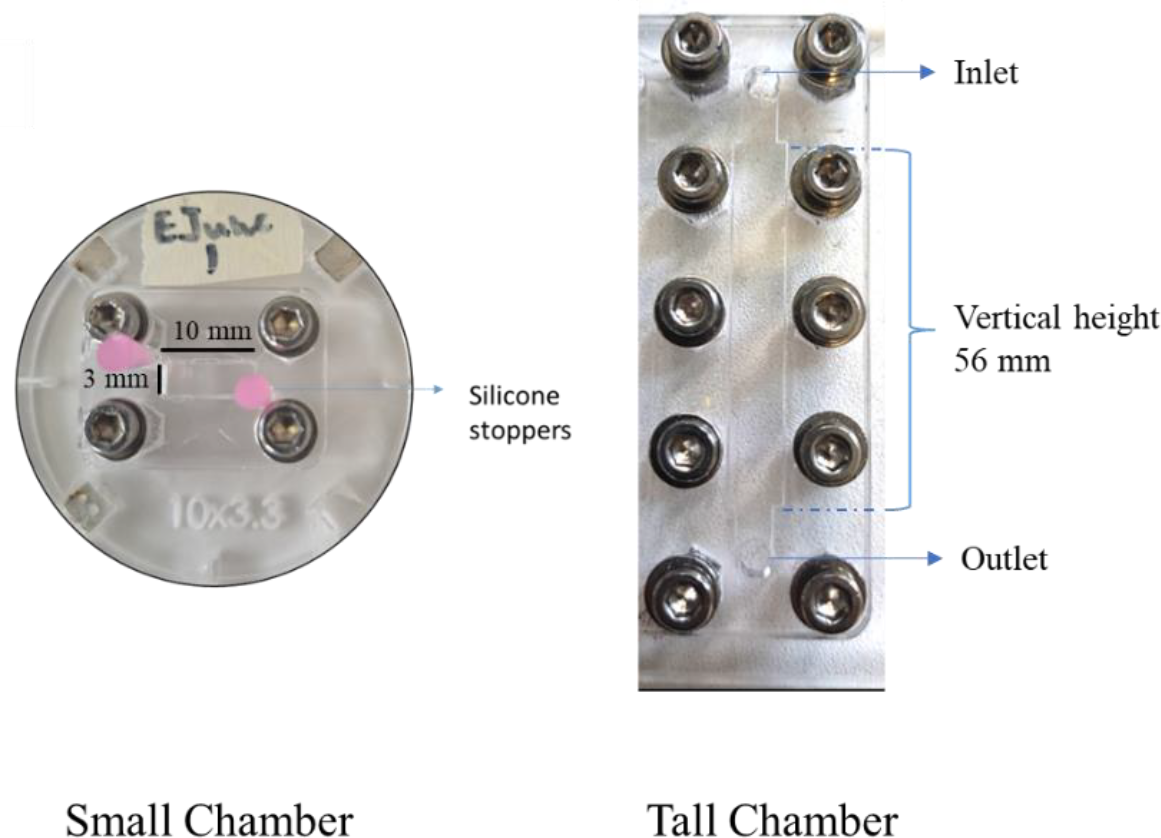

For the tall chamber experiments, a chamber of height 60 mm (shown below) was mounted on a stage which was made by coupling two *xz* translation stages in a series, allowing an increased displacement span of at least 55 mm along the vertical. Cells were filled and allowed 20 minutes to reach a steady distribution (called pre-flip distribution). At ∼ 1.4 x zoom, our camera could capture ∼ 4mm per frame, so we had to move our chamber about 14 times for the camera to capture the entire 56 mm. We excluded the top and bottom 4 mm of our chamber as these regions are close to the inlet and outlet ports and i) imaging is not clear due to projection of stopper (used for sealing the ports) on the imaging plane and ii) avoid any bias arising due to any potential air-medium interface at the ports. For the remaining 56 mm, we stitched the regions together and obtained the complete distribution of the chamber and used for downstream calculations. Scanning the entire chamber (14 regions) took ∼2 minutes, which is less than time taken by cells to cover a region.

### ii) Computing upward bias index

To quantify the upward bias, we used the equation *r* = (*f↑ − f↓*)/(*f↑* + *f↓*), where *f↑* and *f↓* are the concentration of cells in the top and bottom 660 μm of the chamber, respectively. The *r* value lies between -1 to 1. An upward bias value of ∼ -1 means all cells are present in the bottom portion of the chamber, while a value of ∼ +1 means all cells at top. An upward bias value of 0 means a 50:50 distribution of cells between the top and bottom regions.

### iii) Computation of motility parameters from single cell tracking

Cells were put in the small chamber as described above and allowed 20 min. to reach a steady distribution as mentioned above. Then the chamber was rotated by 180 degrees (in 3 seconds) via an Arduino controlled automatic stepper motor (programmed in-house). As the cells started to swim back to their steady state, short videos of about 160 frames @ 16fps (frames per second) were captured. Starting from the first second of swimming (16^th^ frame), 120 frames were chosen, corresponding to 7.5 seconds of swimming. Individual cell coordinates were determined via binarizing and thresholding using the OpenCV2 module in python. Cell coordinates were linked for subsequent frames via the Trackpy module, to obtain a list of all trajectories. From this, speed, horizontal component, and vertical component of swimming velocity was computed. To ensure selection of the best trajectories, a) all tracks shorter than 45 frames and b) tracks with displacement less than ∼ 2 μm (1/10^th^ of a cell body length) and c) speeds < 10 μm /sec or > 200 μm/sec were rejected.

### iv) Computing cell stability from reorientation

Cell steady distribution was reached, and the small chamber was rotated as described above. As the cells reached the middle of the chamber, 3 180-degree flips were applied (with waiting times of 15, 12, and 12 seconds). Out of this, data from the 1^st^ and 2^nd^ flips were used for analysis. Each of these data is termed a ‘sub-replicate (SR)’. Videos were acquired at 16 fps and cells were tracked as described above. 160 corresponding to 10 sec of swimming were sufficient to capture multiple cell reorientation events in each flip. Trajectories less than 45 frames or a net displacement < 20 μm were filtered out. The remaining trajectories were interpolated quadratically (smoothing). For the single trajectory, angular velocity (ω) was obtained for each consecutive frame as a function of the instantaneous angular position (θ). Angular velocities were averaged for a given θ value (ranging from *−*180° to 180°, binned at intervals of 30°). Any ω value greater than 0.5 rads/sec or less than *−*0.5 rads/sec was eliminated. Obtained ω values were fitted to a sinusoidal curve of the form *A* × sin(*x*), and the reorientation time scale (τ_r_) was then obtained as 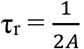.

### v) Measuring the nutrient concentrations

Concentration of two primary nutrients pertaining to phytoplankton growth, nitrate (NO_3_^−^) and phosphate 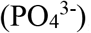, were measured through photometry using Prove600 Spectroquant (Merck) spectrophotometer. For this, cell filtrate was collected from top and bottom of the culture. NO_3_^−^ concentration was analyzed with Nitrate Cell Test in Seawater (method: photometric 0.4 to 13.3 mg/liter NO_3_^−^ Spectroquant) with a minimum detection limit of 6.45 μM. 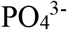 concentration was analyzed with phosphate cellt est (method: photometric 0.2 to 15.3 mg/liter 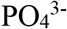 Spectroquant) with a minimum detection limit of 2 μM). Below this limit, values below the respective detection limits are taken as zero.

### vi) Measuring photophysiology using Pulse-amplitude modulated (PAM) chlorophyll fluorometry

A Multiple Excitation Wavelength Chlorophyll Fluorometer (Multi-Color-PAM; Heinz Walz GmbH, Effeltrich, Germany) was used to quantify the maximum photosynthetic quantum 715 yield (*F*_v_*/F*_m_), the maximum electron transport rate (*ETR*_max_), and non-photochemical quenching (*NPQ*) of a cell suspension. For measurements, 1.2 ml cell suspensions were placed into a quartz-silica cuvette (Hellma absorption cuvettes; spectral range, 200-2500 nm; pathlength, 10 mm). PAMWin software saturation pulse (SP) and light curve methods were used to quantify *F*_v_*/F*_m_, *ETR*_max_ and *NPQ* per experiment. The samples were dark adapted (*DA*) for < 1 min. before measurement. Literature suggests that PAM samples should have a DA time of at least 15 min to remove any pre-stress/ difference in samples due to different light/growth conditions present in samples that can bias the measurements. However, i) our samples are all grown and maintained under same conditions and if there is any difference in stress levels, it is possibly due to difference in physiology or other factors between the subpopulations and ii) in our case we are interested to measure the ‘native states’ of the samples i.e. how they are in the culture itself and the longer waiting time, the more identical they start to become. This is also seen in our experiments with *DA* of 20 min. (data not shown), which indicates that in coexistence, the stress levels (*NPQ)* are different than in isolation, similar to swimming behaviors of isolated vs mixed samples.

### vii) Measuring pH of the spent media

For measuring pH of medium/samples, ∼3ml pure medium (kept in incubator under same condition as cells)/ spent media were collected into a 15ml plastic falcon tube. Into this, the electrode of a Mettler Toledo FiveEasy Plus (FP20) Benchtop pH/mV Meter (post-calibration with buffers of pH 4.01, 7.0 and 9.21) was inserted and pH values were obtained.

**Table 1.**
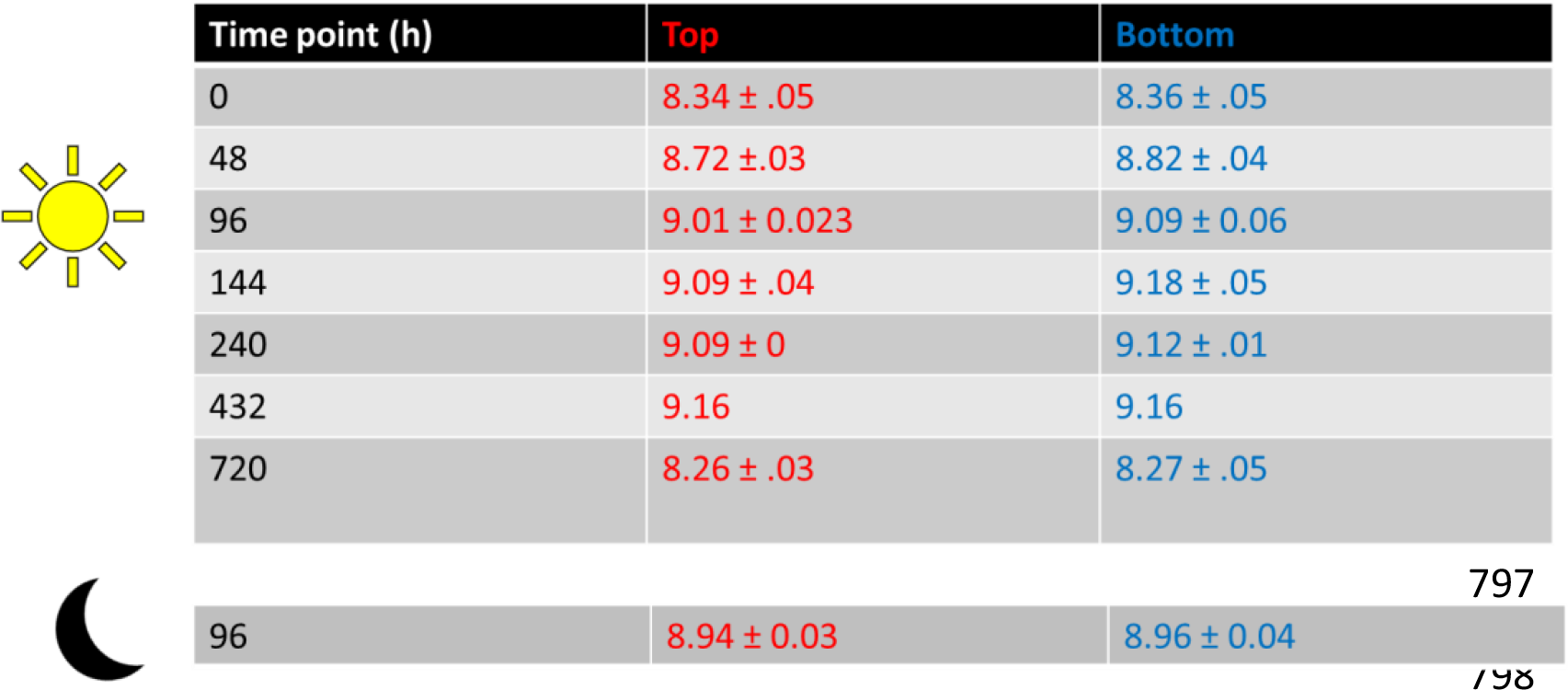
pH of the samples measured in experiments.

| Time point (h) | Top | Bottom |
| --- | --- | --- |
| 0 | 8.34 ± .05 | 8.36 ± .05 |
| 48 | 8.72 ± .03 | 8.82 ± .04 |
| 96 | 9.01 ± 0.023 | 9.09 ± 0.06 |
| 144 | 9.09 ± .04 | 9.18 ± .05 |
| 240 | 9.09 ± 0 | 9.12 ± .01 |
| 432 | 9.16 | 9.16 |
| 720 | 8.26 ± .03 | 8.27 ± .05 |
| 96 | 8.94 ± 0.03 | 8.96 ± 0.04 |

## Single cell experiment

### i) Computing fore-aft asymmetry parameters from cellular morphology

To observe morphology as a function of time, 500 μl cells were pipetted out from a culture tube and put into 1.5 ml glass vials. This was covered with a parafilm and perforated with a single needle and stored in the same incubator as the culture tube (22° C). 10 μl cells were pipetted out at 0 (immediately after cells were collected from the culture tube), 20, 60 and 120 min. These were put on a glass slide and covered with a coverslip and put under a microscope (Olympus CKX53 inverted) and viewed in phase contrast mode using a camera (same as above). Short videos of cells were captured at 82 fps and at least 10 cells were imaged per time point.

To analyze morphology, a single frame showing a cell in maximum focus was chosen from a video manually. This was then cropped around the cell and using OpenCv2 module, points were drawn along the cell’s external contour and the *xy* coordinates per point were generated.

These were converted to polar co-ordinates and fit to a three-parameter 3D spheroid surface **[55]** which, in 2D, translates to:

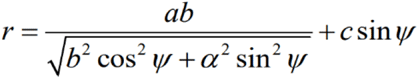

i.e., an ellipsoid with *a* and *b* as major, minor semi-axis **[7]**. This ellipsoid is fore-aft asymmetric i.e., in the direction of the major axis and this asymmetry is captured by the parameter *c*.

### ii) Computing fore-aft (a)symmetry parameters from cellular morphology in presence of the spent media

1.4 ml cell suspension of one type (top or bottom) was collected and 800 μl of this suspension were put into two 1.5 ml glass vials (400 μl each). The rest was filtered using a 1 μm Whatman^®^ Puradisc 25 syringe filter, to form the filtrate (as described previously) of the same type. Next, 1ml of the other cell type was collected immediately and passed through the filter to form the opposite supernatant. 150 μl of both supernatants were collected individually and put into one of the vials to form a total of two experimental cases per sample type. For each experiment, 10 cells were imaged under the same microscope as above, at least at 50 fps and analyzed using the same protocol as described above.

### iii) Cell mechanics model and calculation of the stability diagrams

To determine the propensity of the cells to maintain their swimming stability (and their inherent cell reorientation timescale) under diverse condition, we employ a cell-level mechanistic formulation. The model takes into account the various forces and torques a phytoplankton cell experiences by virtue of its motion and cytoplasmic organelle (nucleus) position. Our primary aim is to identify the dominant forces and torques a motile cell encounter and establish the factors that determine their up-swimming stability. A cell generates a propulsive force, ***P*** (because of its flagellar dynamics) to maintain its active motion. The weight of the cell (due to combined influence of the nucleus, lipids and cytoplasm) and the upthrust (attributed to the finite cellular volume) on the cell are inherent forces owing to its physiological configuration. In addition, the cell motion induces a drag force attributed to the cell geometry and viscous effects. Torques on the cell structure is estimated about its centroid (or center of buoyancy, *C*_*B*_, see Fig. 6 below). The torque contributions on the cell mechanics are the following: torque due to nucleus and lipids (when their center of residence is not on *C*_*B*_), torque originating from viscous drag (in case of asymmetric cellular geometry), and resistive (viscous) torque due to cell rotation (with rotation speed *ω*). With the above physical considerations, we balance the forces and torques in each component to formulate the reduced-order model that reads (see Fig. 6):

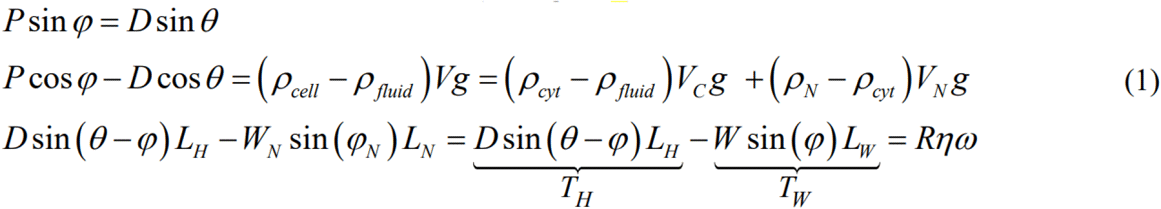

**Fig. 6.**
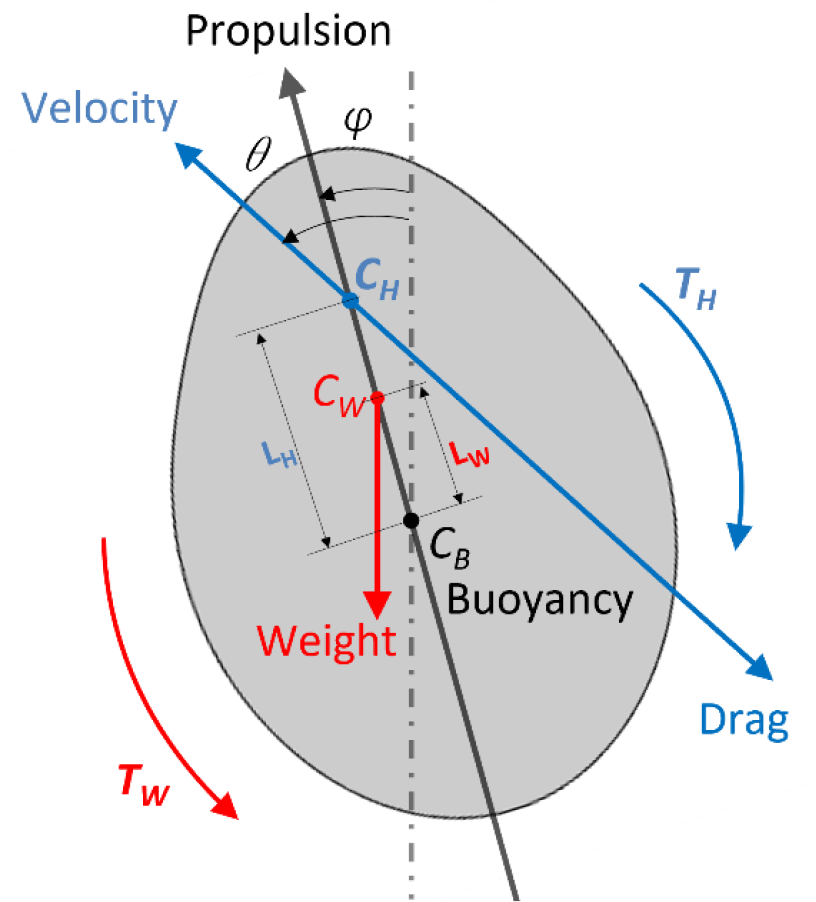
Schematic of the cell geometry showing the forces and torques (about the center of buoyancy, *C*_*B*_) acting on a phytoplankton cell.

The symbols *ρ, V, W, η* and *L* denotes the density, volume, weight, medium viscosity, and distance from cell centroid respectively. Some of the symbols carry the subscripts *cyt, fluid, C, H*, and *N* which, respectively, refers to the cytoplasm, background medium (within which the cell remains submerged), the hydrodynamic center of the cell, and the cell nucleus. *φ* is the angle between the line of action of the propulsive force, ***P*** (attributed to the resultant flagellar motion) and the line of action of gravity. Here *φ* and P are unknowns, which needs to be determined as part of the solution. In reality, the motion of the cell does not follow the line of action of ***P***, hence an angular offset *θ* (an experimentally observable parameter) with the vertical is assumed along which the cell moves. *φ*_*N*_ is the angle between the direction of gravity (vertical line) and the line joining *C*_N_ and *C*_B_ (note *φ* = *φ*_*N*_, since we assume the center of gravity of the nucleus to lie on the major axis). *D* denotes the drag force whose knowledge requires the detail of the cellular geometry and its interaction to the surrounding fluid, the details of which are provided below.

We describe the axisymmetric cell geometry with the generic equation 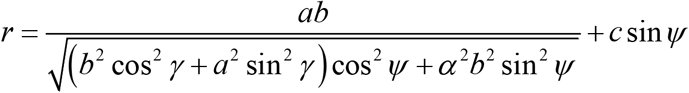 where the symbols *a, b* (*a>b*), *ψ* (*−π*/2 *<ψ < π*/2), and *γ* (0 *< γ <* 2*π*) represents the major axis length, minor axis length (equal to the semi-major axis length), polar angle, and azimuth angle, respectively. Here *c* implies the deviation from the symmetric shape along the major axis (fore–aft direction) and *r* denotes the position vector of the points on the cell surface (from the origin) as a function of the polar and azimuth angles.

The fore-aft asymmetry (value of *c*) is quantified using the phase-contrast microscopy images of the cells whose contours are fitted with Equation 1 and *γ* = 0, resulting in the form 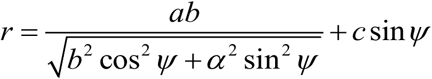. Note that for a symmetric cell geometry (*c* = *0*), the hydrodynamic center (*C*_H_) falls on the cell centroid (*C*_B_), and *L*_*H*_ vanishes. With the consideration that the cell shape may be assumed as a prolate spheroid, the drag of a symmetric prolate ellipsoid is expressed as *D*_P,*⊥*_ = 6*πηr*_*eq*_*UK*_P,*⊥*_ where *U* and *K* are the translational velocity and the shape factor, respectively, while P (_*⊥*_) denotes the parallel (perpendicular) direction with respect to the major axis. The shape factors have the form 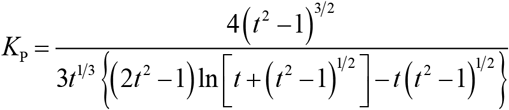 and 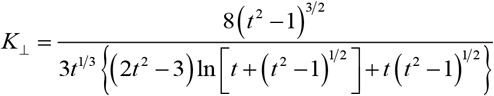for prolate spheroids **[56, 57]** where *t=a/b*. The net drag on the cell is dictated by its orientation and is given by *D* = *D*_P_ cos(*α*) *+ D*_*⊥*_ sin (*α*) (*D*_P_ and *D*_*⊥*_ are the drag forces parallel and perpendicular to the major axis of the cell shape, respectively, and *α* = *θ −φ*).

*R* represent the coefficient of hydrodynamic rotational resistance and has the form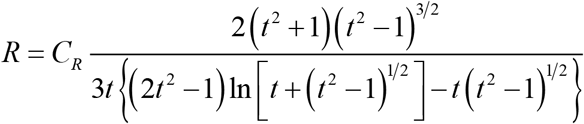 **[56]** where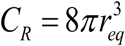. With *R* defined, the viscous torque on a prolate spheroid is estimated using *τ* = *Rηω* where *ω* is the angular rotation rate (rad/s). Our aim is to obtain the angular rotation rate ***ω*** from the above set of three coupled equations (Equation 1). Using the experimentally known values (Table 2) we draw a stability phase-plot that presents the value of the angular rotation rate as a function of shape asymmetry and cell geometry (estimated by varying *L*_H_/*a* and *L*_W_/*a*). The stability phase plots demarcate the regions of stable up-swimmers from stable down-swimmers, thereby encompassing the various stability conditions of cell motility as a function of various environmental factors. Here we note that since the nucleus resides above the cell centroid, the effects of the lipids on the cell stability may be safely ignored.

**Table 2.**
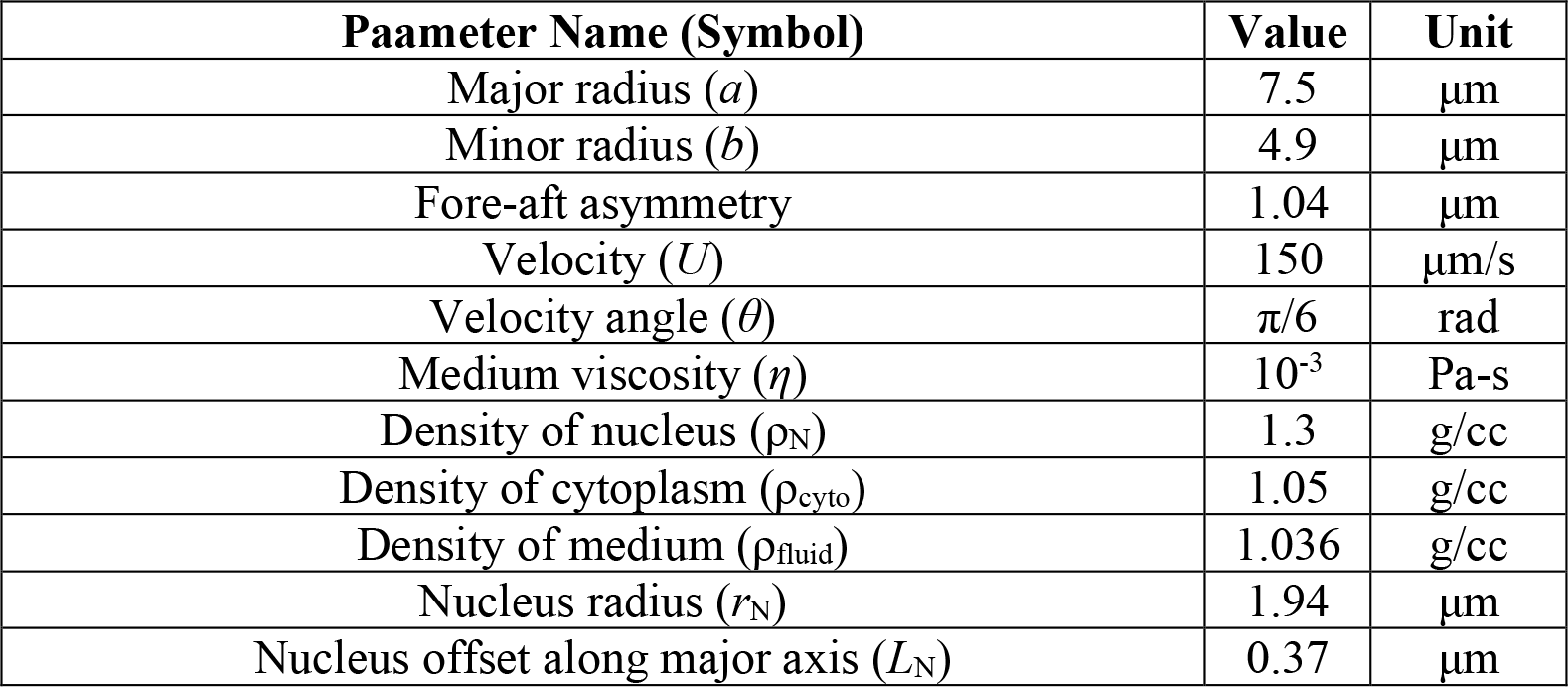
Parameters used to draw the phase-plot diagram for the cell stability.

| Parameter Name (Symbol) | Value | Unit |
| --- | --- | --- |
| Major radius ($a$) | 7.5 | $\mu\text{m}$ |
| Minor radius ($b$) | 4.9 | $\mu\text{m}$ |
| Fore-aft asymmetry | 1.04 | $\mu\text{m}$ |
| Velocity ($U$) | 150 | $\mu\text{m/s}$ |
| Velocity angle ($\theta$) | $\pi/6$ | rad |
| Medium viscosity ($\eta$) | $10^{-3}$ | Pa-s |
| Density of nucleus ($\rho_N$) | 1.3 | g/cc |
| Density of cytoplasm ($\rho_{\text{cyto}}$) | 1.05 | g/cc |
| Density of medium ($\rho_{\text{fluid}}$) | 1.036 | g/cc |
| Nucleus radius ($r_N$) | 1.94 | $\mu\text{m}$ |
| Nucleus offset along major axis ($L_N$) | 0.37 | $\mu\text{m}$ |

## Acknowledgements

The authors thank Robert Himelrick and Nicolas Tournier for the support with fabrication of experimental chambers.

## SUPPLEMENTARY MATERIAL

### This file contains

Supplementary figures and captions for Figures S1 to S18

## Supplementary Figures

**Fig. S1.**
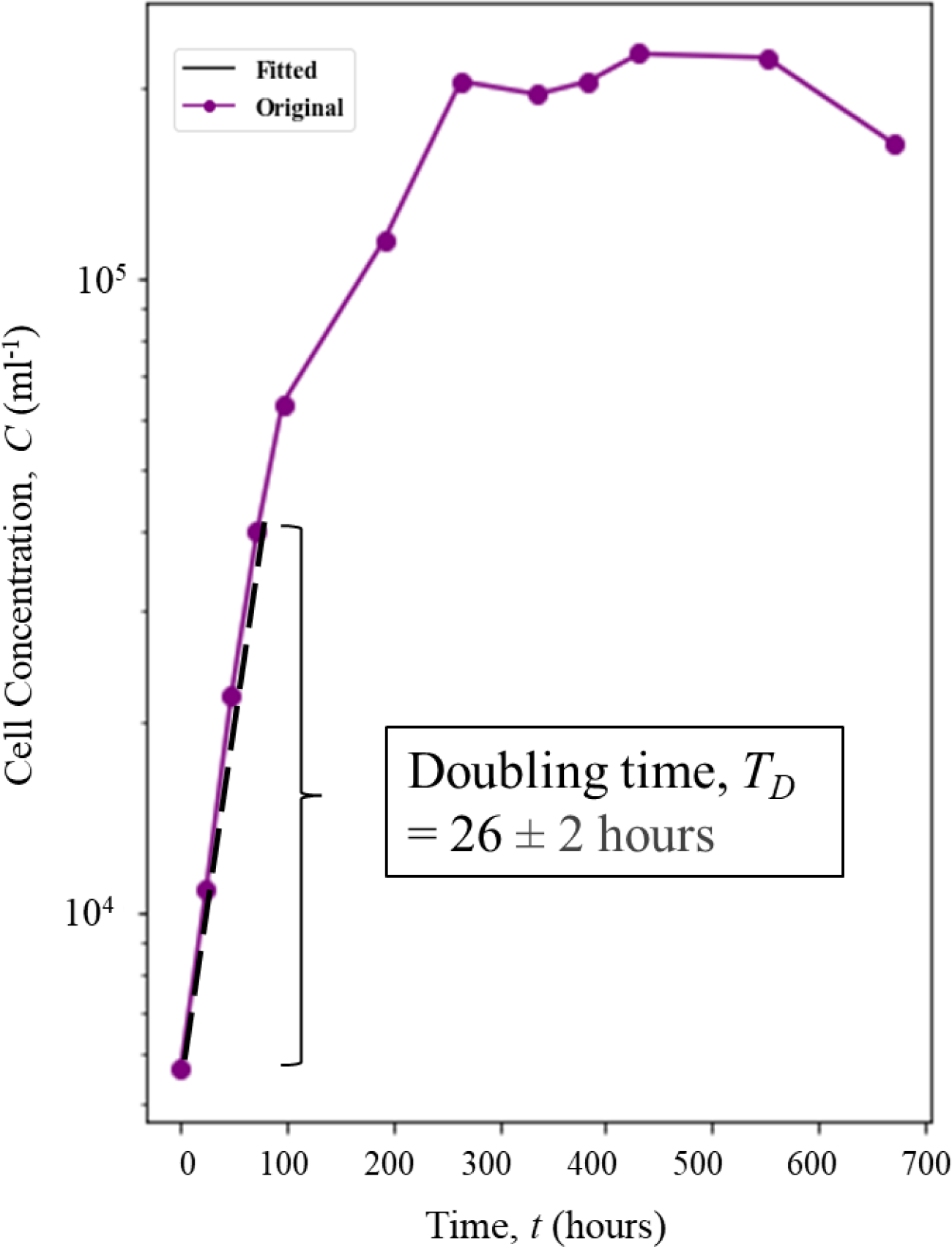
Quantification of doubling time from HA 452 growth curve. In order to compute the doubling time, the growth curve shown in Fig.1 A (main text), was plotted in a logarithmic scale, shown in purple. The exponential phase of growth is fitted to an exponential function (shown in black lines). The coefficient of the exponent gives us the growth rate from which doubling time is calculated (Materials and Methods). Doubling time, *T*_*D*_ ranges from 24 to 28 hours which matches well with ranges of other phytoplankton mentioned in literature (see main text).

**Fig. S2.**
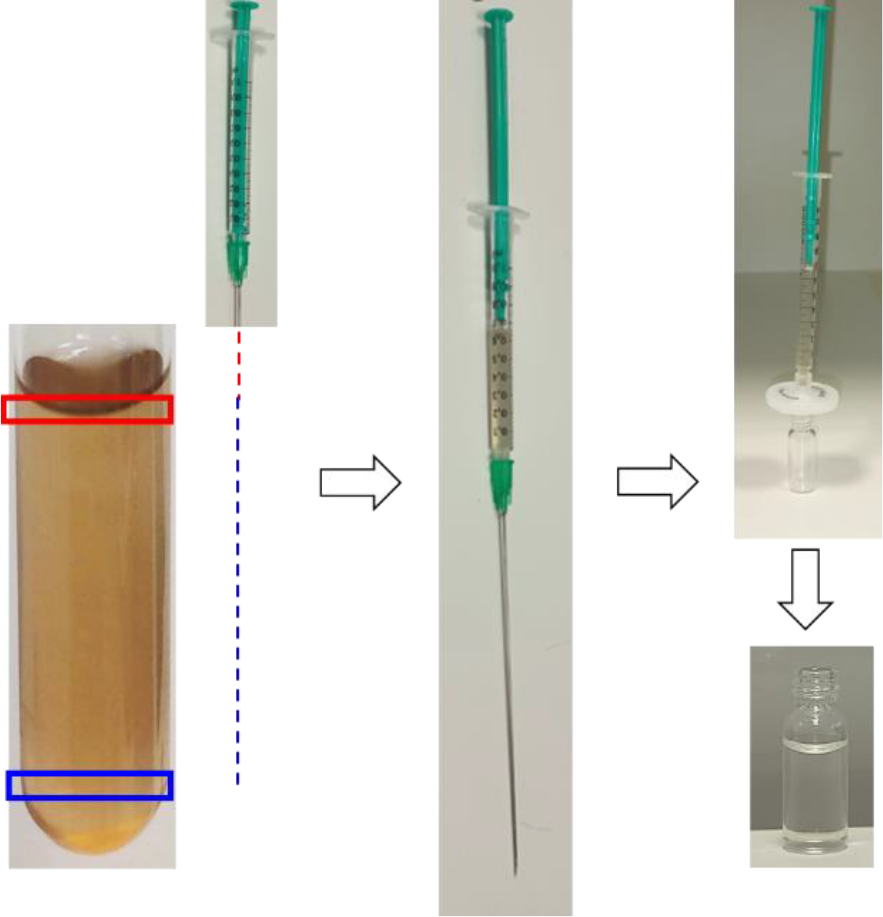
Generation of filtrate from a cell culture. To generate filtrates, cell samples are collected from a growing culture (left most panel) from either the top (red box) or bottom (blue box) region. Collection is done using a needle and syringe (Materials and Methods). The red or blue dotted lines indicate the depth of needle for collection of top or bottom sample. Once samples are collected (middle panel), they are passed through a 1 μm pore sized syringe filter (right panel, top) and collected into a glass vial (right panel, bottom). These are then used for measuring pH, nutrients, and response of cells in presence or absence of same or opposite filtrates.

**Fig. S3.**
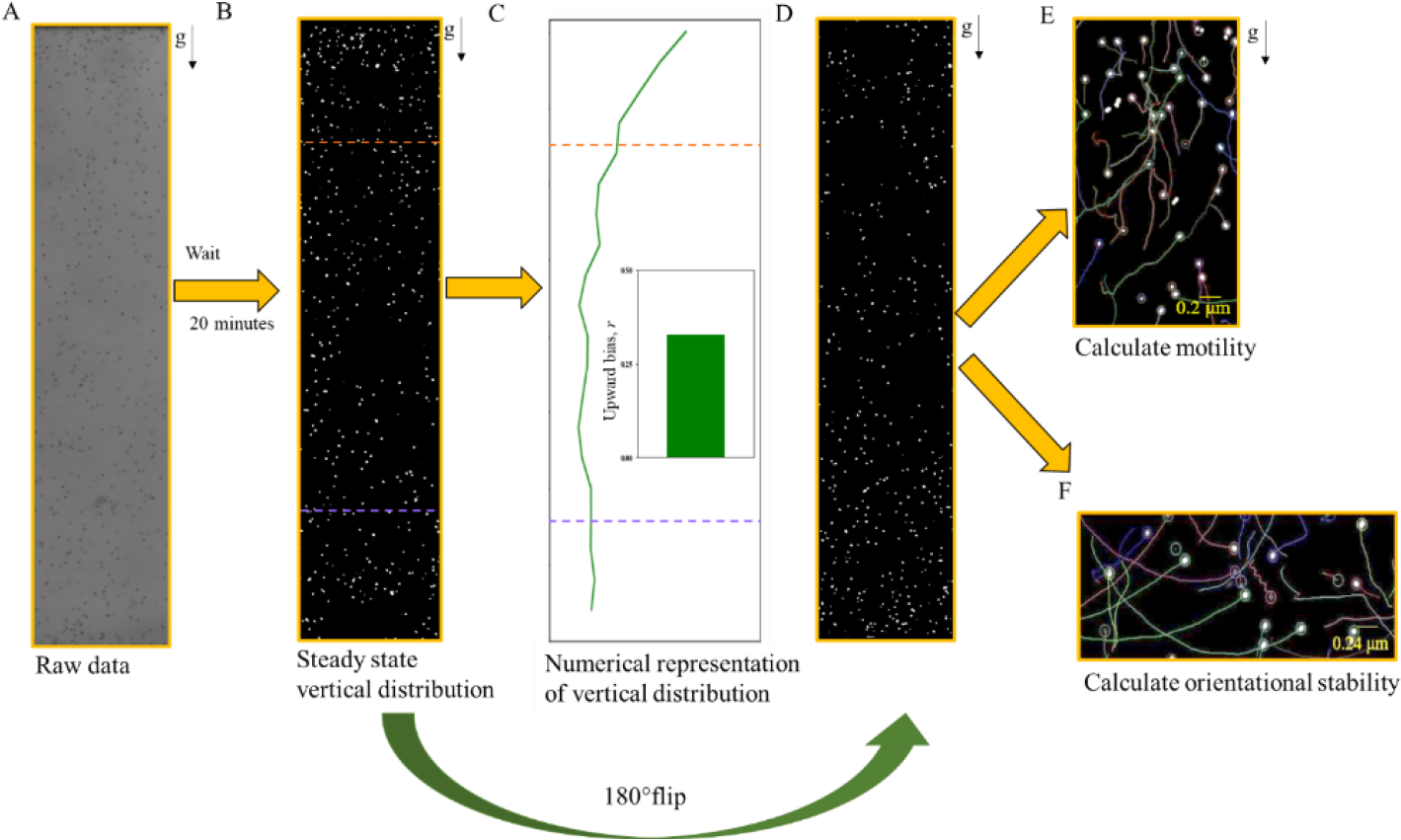
Image analysis pipeline. This figure shows a general pipeline used for analyzing raw data (panel **(A)**) and extract parameters like upward bias, motility and stability. These parameters have been mentioned and used multiple times throughout the paper and uses the same approach as shown here. First one takes the raw data and then binarizes it for downstream analyses. **(B)** shows one such binarized image of cells in a millifluidic chamber, after 20 minutes from the point of insertion. **(C)** shows how the binarized input is used to compute vertical distribution, shown in green line (for details, see Materials and methods). From this, upward bias value is calculated by considering the cell concentration in the top and bottom 1/5^th^ of the chamber, indicated by the dark orange and lavender dotted lines. **(D)** To calculate motility and orientational stability, after an equilibriation time (here 20 minutes), the chamber is flipped by 180 degrees. Cells at the top pre-flip are now at the bottom and start to swim up. **(E)** As cells swim up, a short video is captured which is analysed to obtain swimming trajectories from which motility parameter like speed and swimming direction can be obtained for individual cells. **(F)** Flipping the chamber also changes cellular orientation from their stable configuration and they start to reorient, as shown by the arc-like trajectories.

**Fig. S4.**
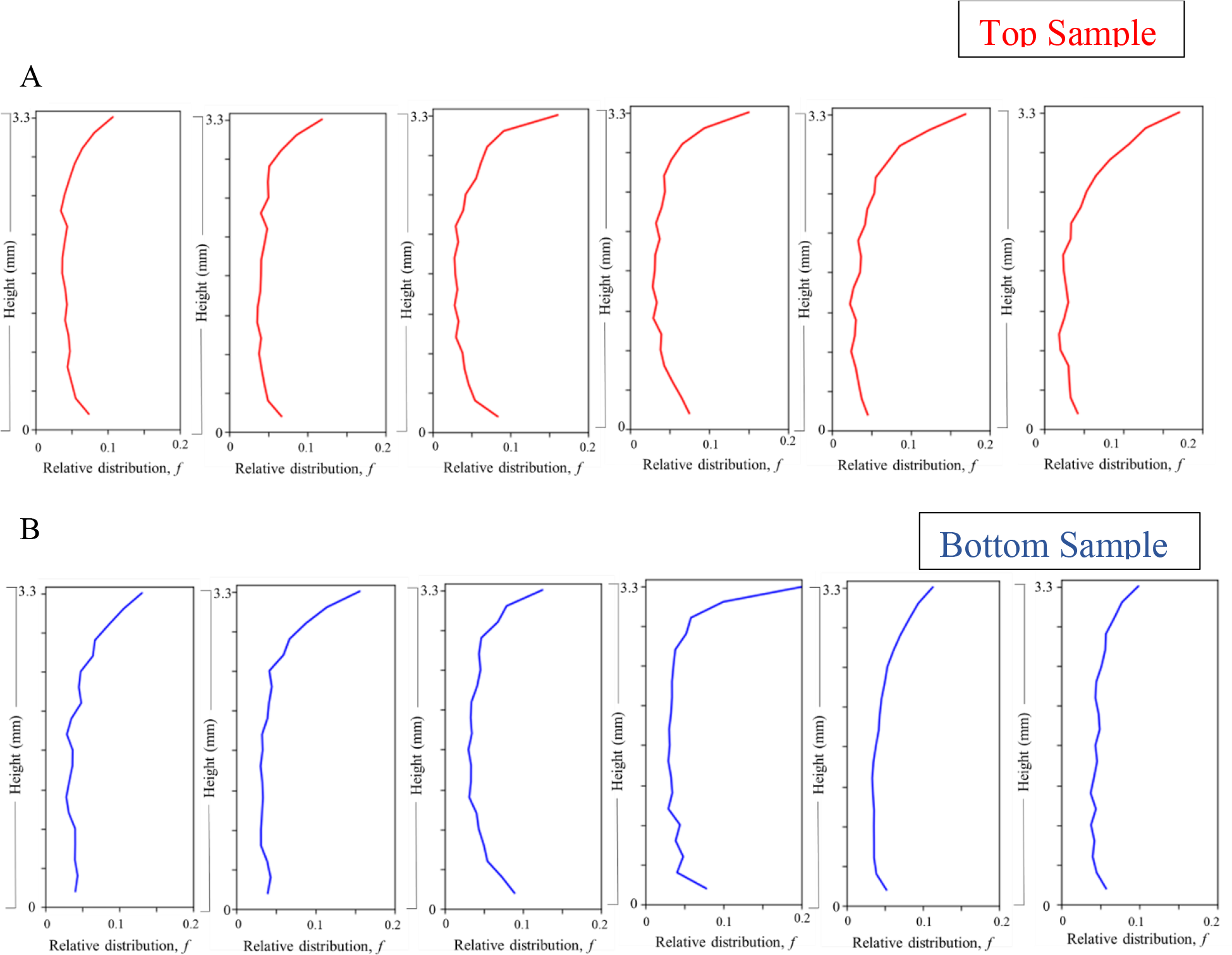
Vertical distribution of cells in a multifluidic chamber. To observe the vertical distribution of cells, samples were collected from top (red) and bottom (blue) and placed into a multifluidic chamber of vertical height 3.3 mm. After 20 minutes, the entire height was imaged for subsequent cell count. **(A)** shows the distribution of top cells with each panel corresponding to one replicate (of 6 replicates, 3 biological replicates each having 2 technical replicates). **(B)** shows the same but for bottom cells. First two panels show technical replicates of the first biological replicate and the rest follows that order.

**Fig. S5.**
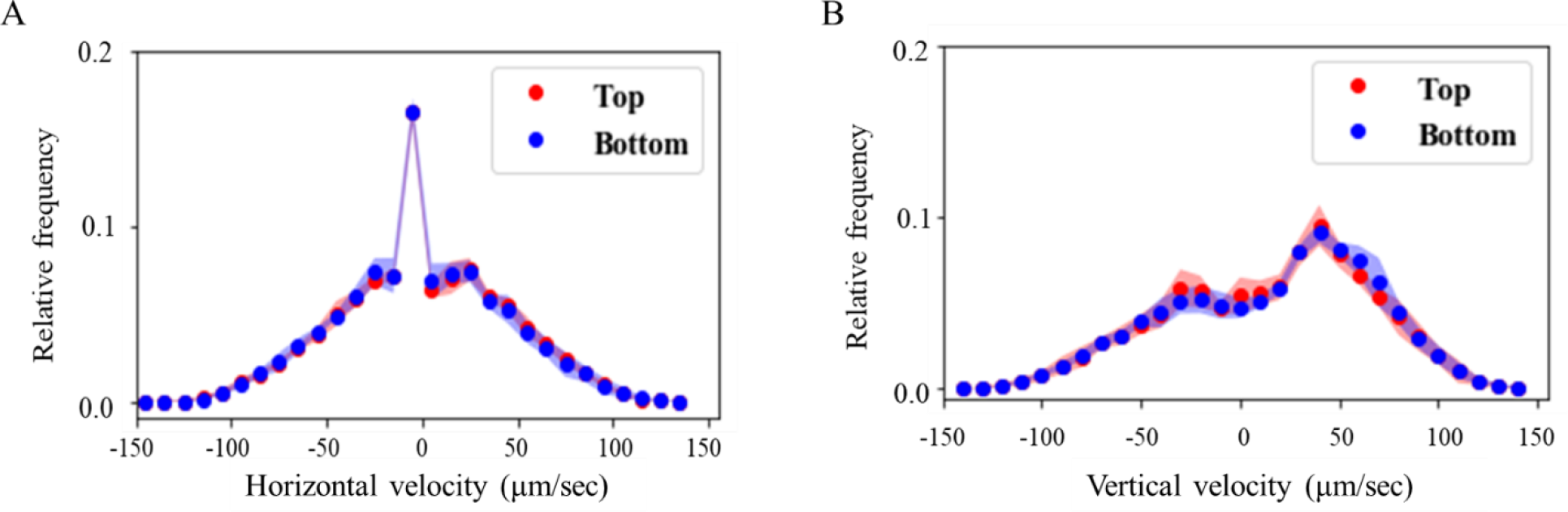
Motility parameters of cells. While Fig. 1 H of main text shows that the mean absolute swimming speed of top and bottom cells are similar, we also wanted to see if their directionality remains similar identical as well (reason in main text). **(A)** shows the relative distribution of mean horizontal swimming speed of all 6 replicates for top and bottom cells (red and blue respectively). Horizontal velocity is divided into two sections about 0. Cells swimming towards their right from their initial position is assigned positive velocity and those swimming left are assigned negative velocities. Results indicate that the majority population of both samples have a mean horizontal speed of 0 μm/s, meaning that on a population scale, cells do not show much displacement in the horizontal direction. **(B)** shows the relative distribution of mean vertical swimming speed of the same. Vertical velocity is divided into two sections about 0. Cells swimming against gravity from their initial position is assigned positive velocity and those swimming towards, are assigned negative velocities. (Direction of gravity, *g*, is shown by the black arrows to the right of both panels) Mean vertical velocity of both samples peak at around 35 μm/s, meaning that the population as a whole move actively against gravity, precisely in the negative vertical direction.

**Fig. S6.**
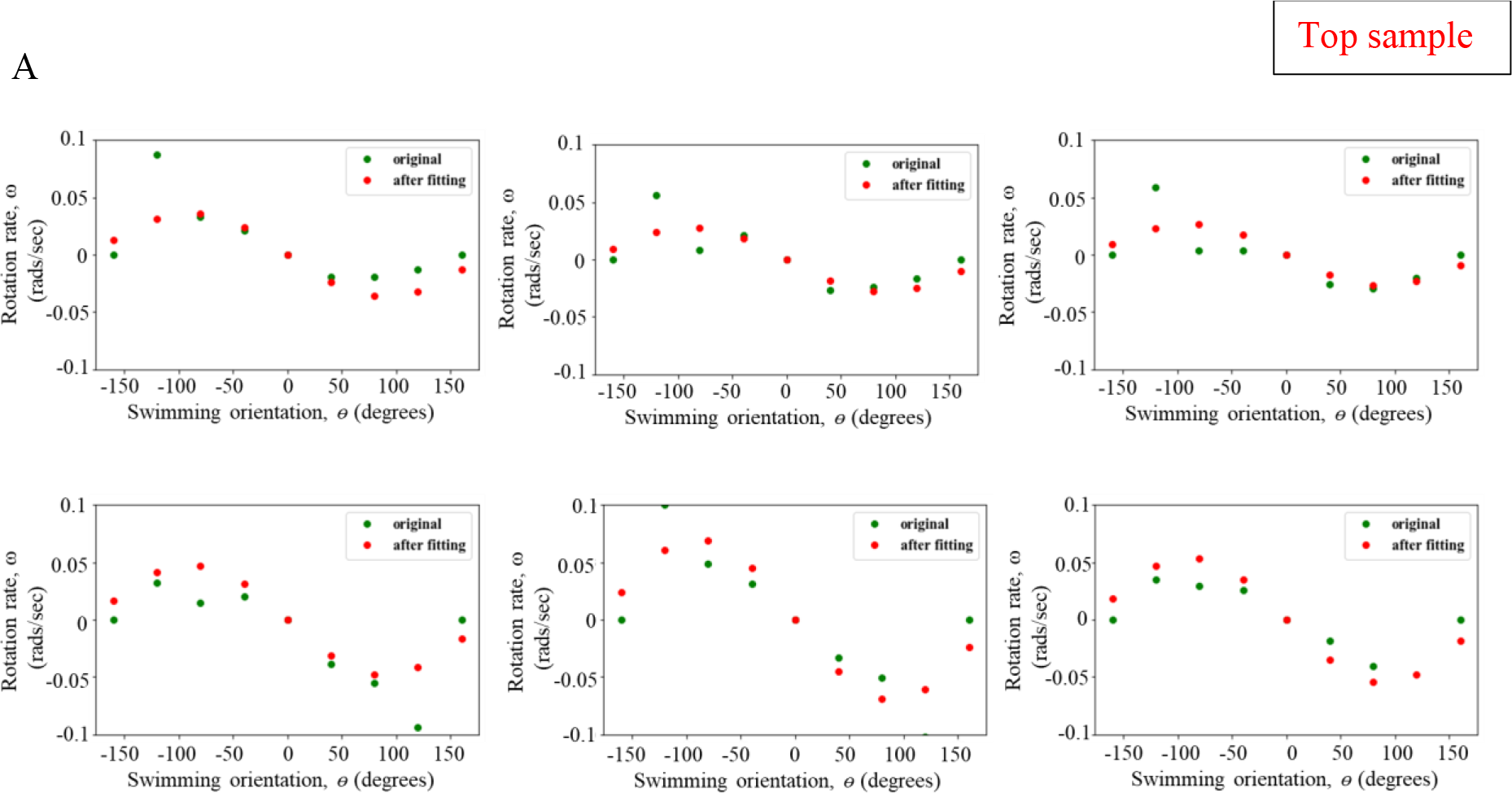
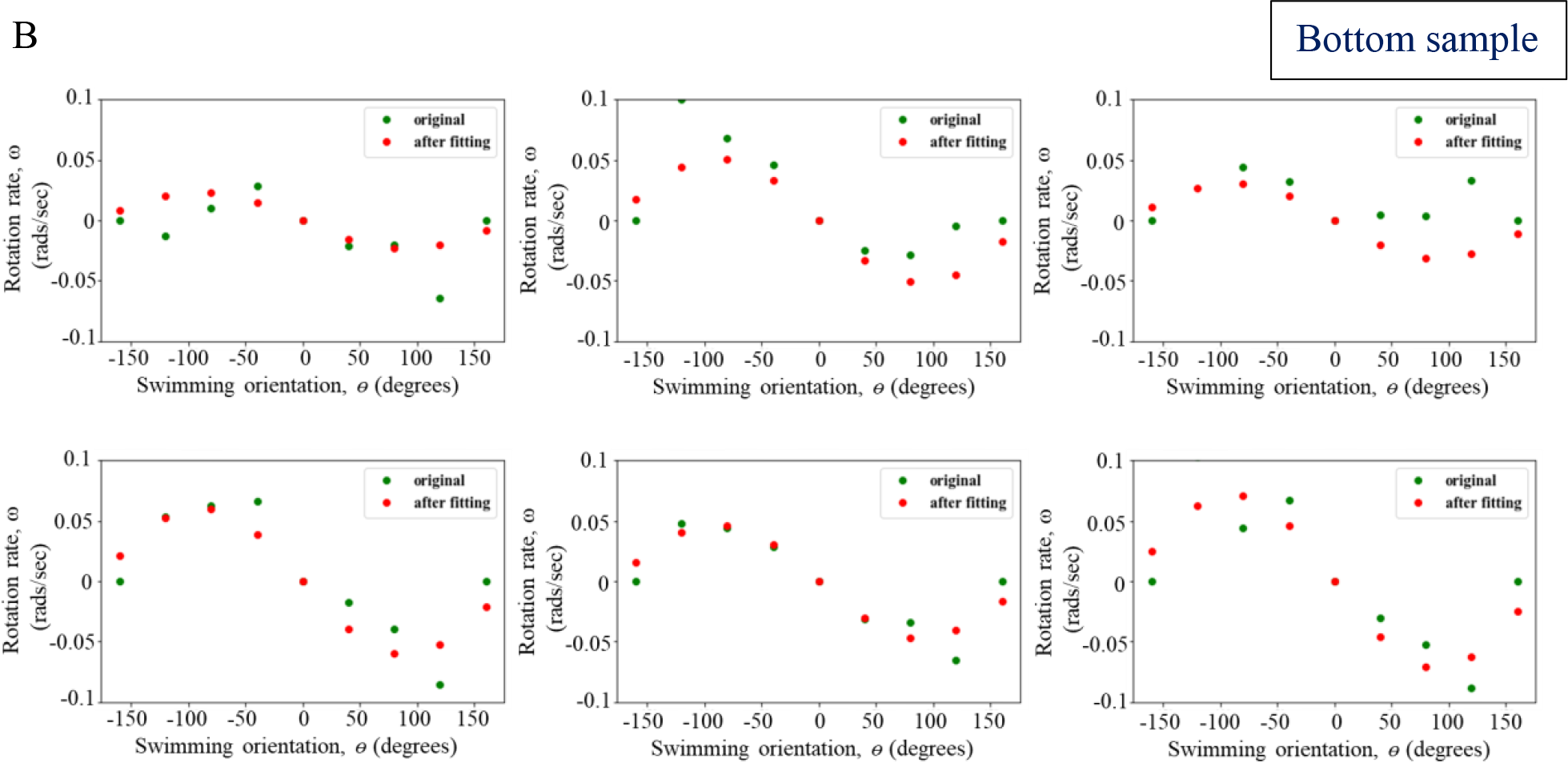
Quantification of the orientational stability of cell samples. To compute the stability of top and bottom samples, rotation rate (ω) of cells were plotted as a function of the instantaneous angular direction (θ). The time of experiment was day (11 hours). **(A)** shows the stability of top cells with each panel corresponding to one replicate. Red dots correspond to experimental data and green dots to the sinusoidal curve obtained by fitting a sinusoid to the experimental data. (B) shows the same but for bottom cells. The amplitude, *A* of the fitted curve was used to obtain the reorientation timescale, *B* as 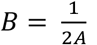. Note that there are 12 replicates: 3 biological replicates x 2 technical replicates x 2 subreplicates, where 1 subreplicate is obtained by imaging 1 technical replicate for two consecutive flips, as detailed in the Materials and methods section. Here the subreplicates per technical replicate has been combined together, thus showing a total of 3x2x1= 6 plots.

**Fig. S7.**
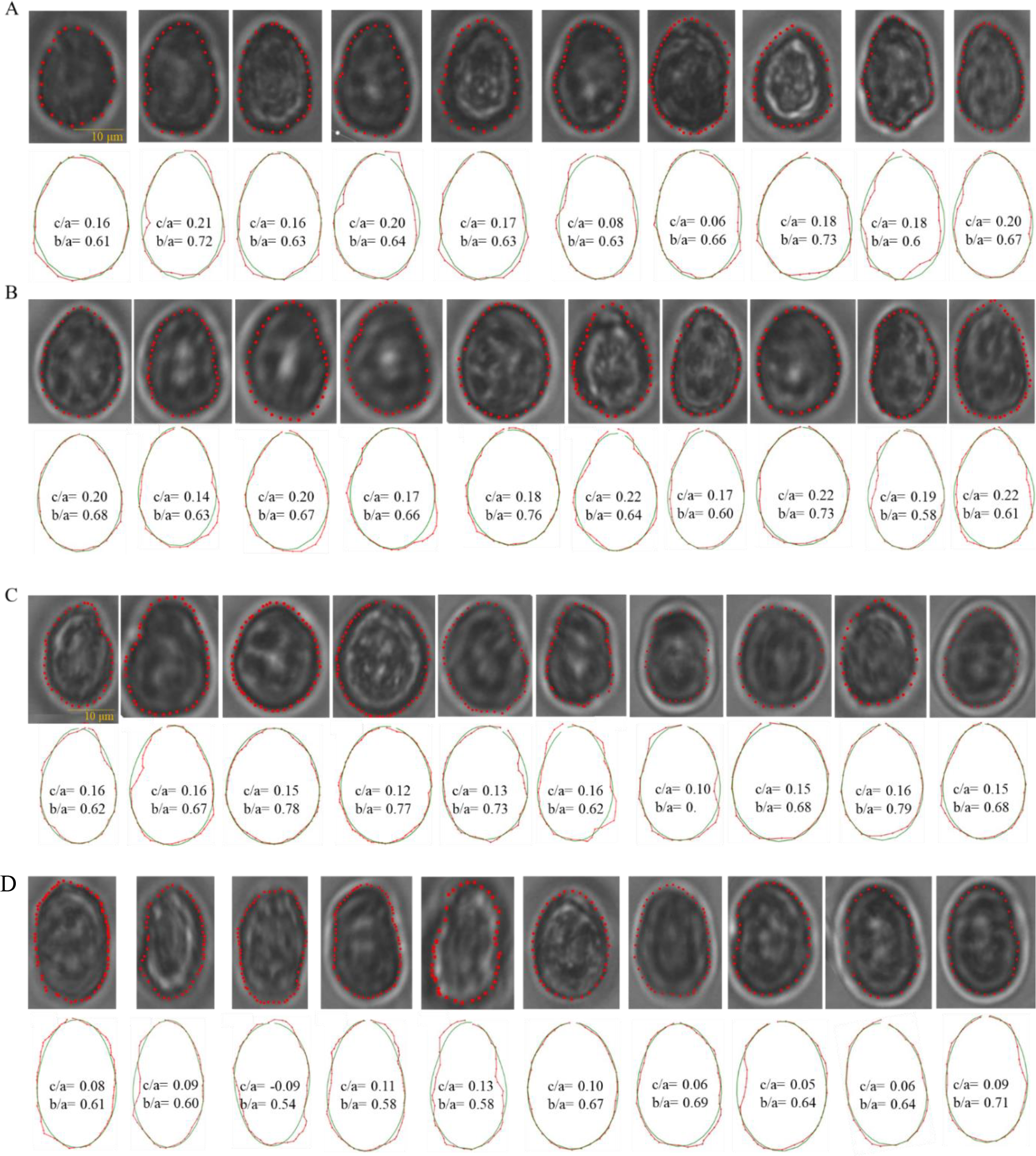
Quantification of morphological parameters of top cells. The top row of each panel shows phase contrast images of top cells (*N* = 10). The external contour of each cell was marked, and co-ordinates were extracted. These were then fed into an image analysis pipeline to fit each contour with a fore-aft asymmetric ellipse. The *c/a* parameter is a quantification of this asymmetry. Panels A to D represents distinct time points of 0, 20, 60 and 120 min.

**Fig. S8.**
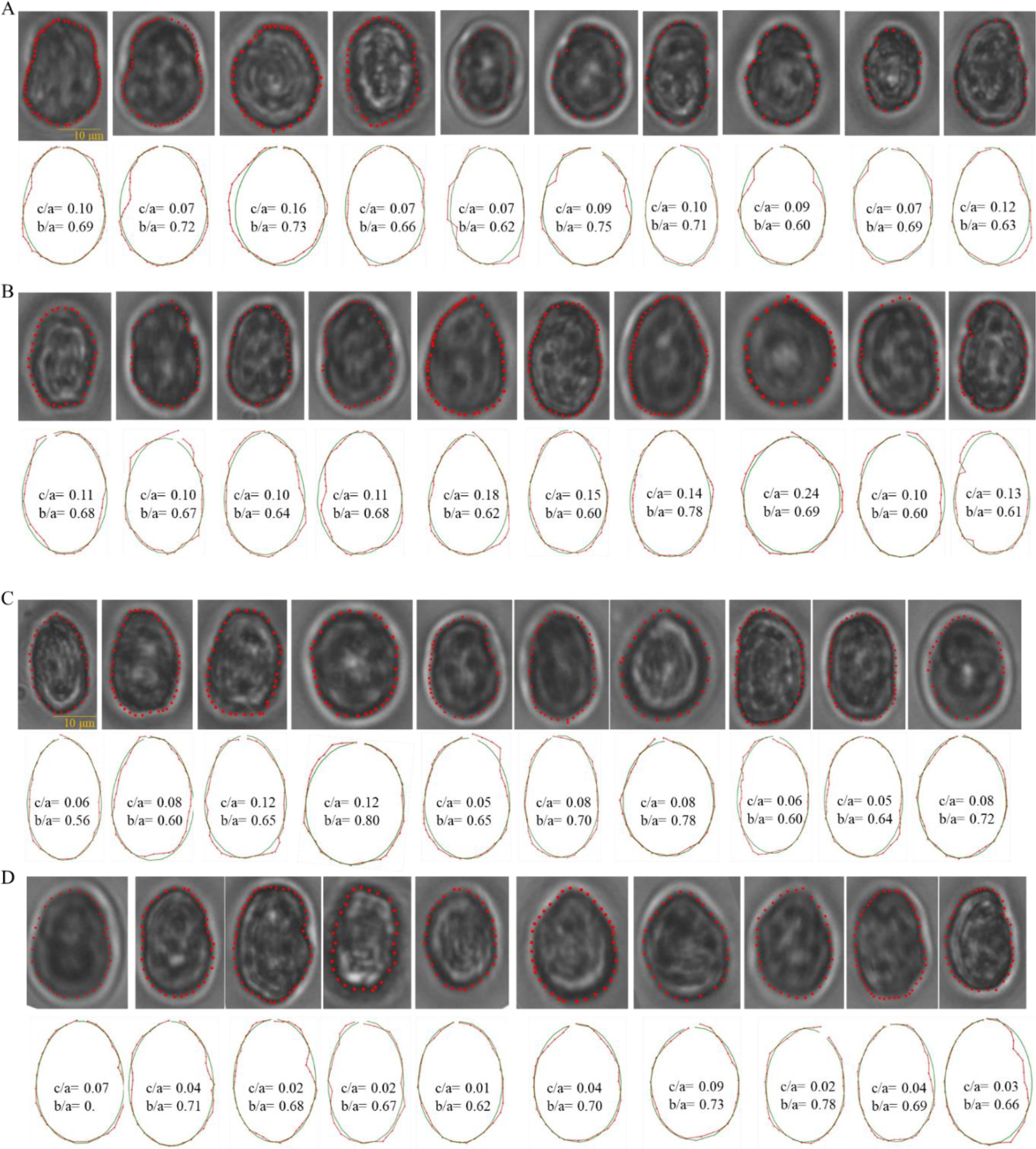
Quantification of morphological parameters of bottom cells. The top row of each panel shows phase contrast images of bottom cells (*N* = 10). The external contour of each cell was marked, and co-ordinates were extracted. These were then fed into an image analysis pipeline to fit each contour with a fore-aft asymmetric ellipse. The *c/a* parameter is a quantification of this asymmetry. Panels A to D represents distinct time points of 0, 20, 60 and 120 min.

**Fig. S9.**
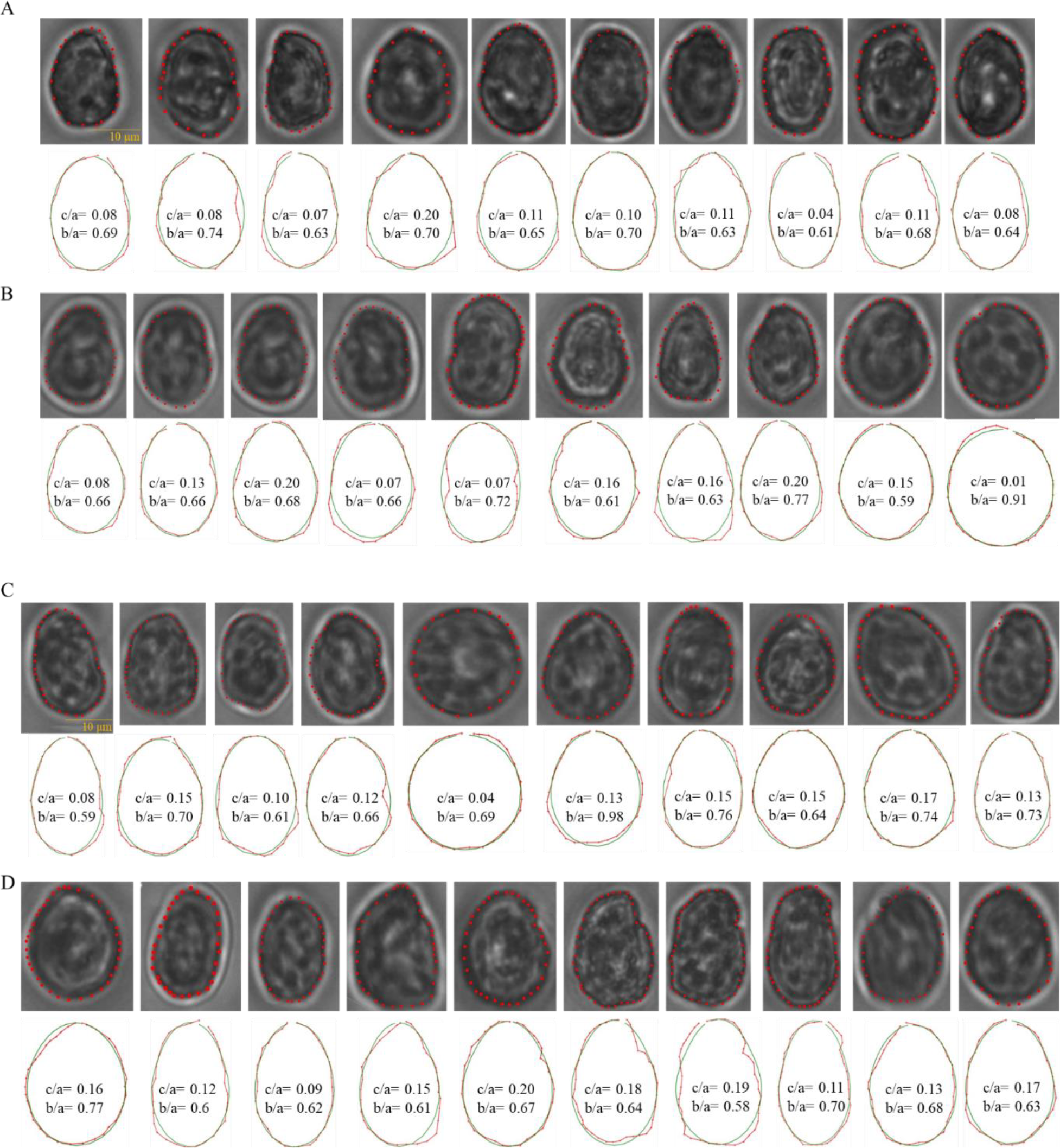
Quantification of morphological parameters of mixed cells. The top row of each panel shows phase contrast images of 50:50 mixed cells (*N* = 10). The external contour of each cell was marked, and co-ordinates were extracted. These were then fed into an image analysis pipeline to fit each contour with a fore-aft asymmetric ellipse. The *c/a* parameter is a quantification of this asymmetry. Panels A to D represents distinct time points of 0, 20, 60 and 120 min.

**Fig. S10.**
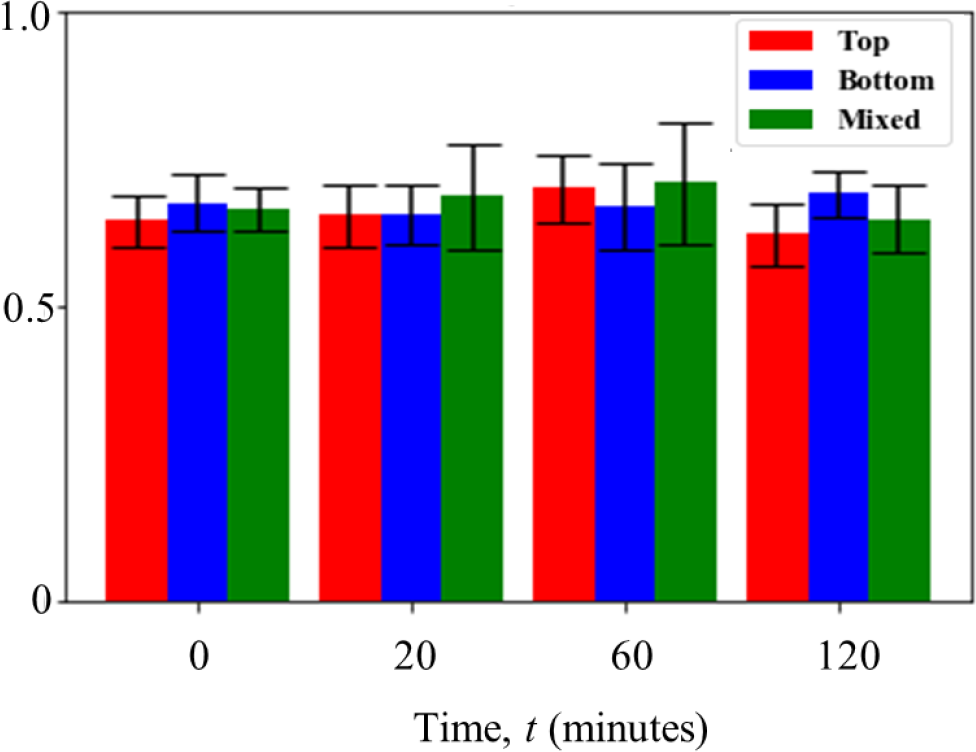
*b/a* values of cells over time. Complementing the *c/a* values shown in main text, this figure shows the *b/a* values, the ratio of the minor to major semi axis, that gives an idea of the shape of cells. A value of 0 means cells are highly asymmetric, while that of 1 means highly rounded cells. For top cells (red bar), the mean (± s.d.) values over 0, 20, 60 and 120 minutes are 0.65 ± 0.04, 0.66 ± 0.05, 0.7 ± 0.06 and 0.62 ± 0.05. For bottom cells (blue bar), they are 0.68 ± 0.05, 0.66 ± 0.05, 0.67 ± 0.07 and 0.7 ± 0.04. For mixed cells (green bar), they are 0.66 ± 0.04, 0.69 ± 0.09, 0.7 ± 0.1 and 0.65 ± 0.06.

**Fig. S11.**
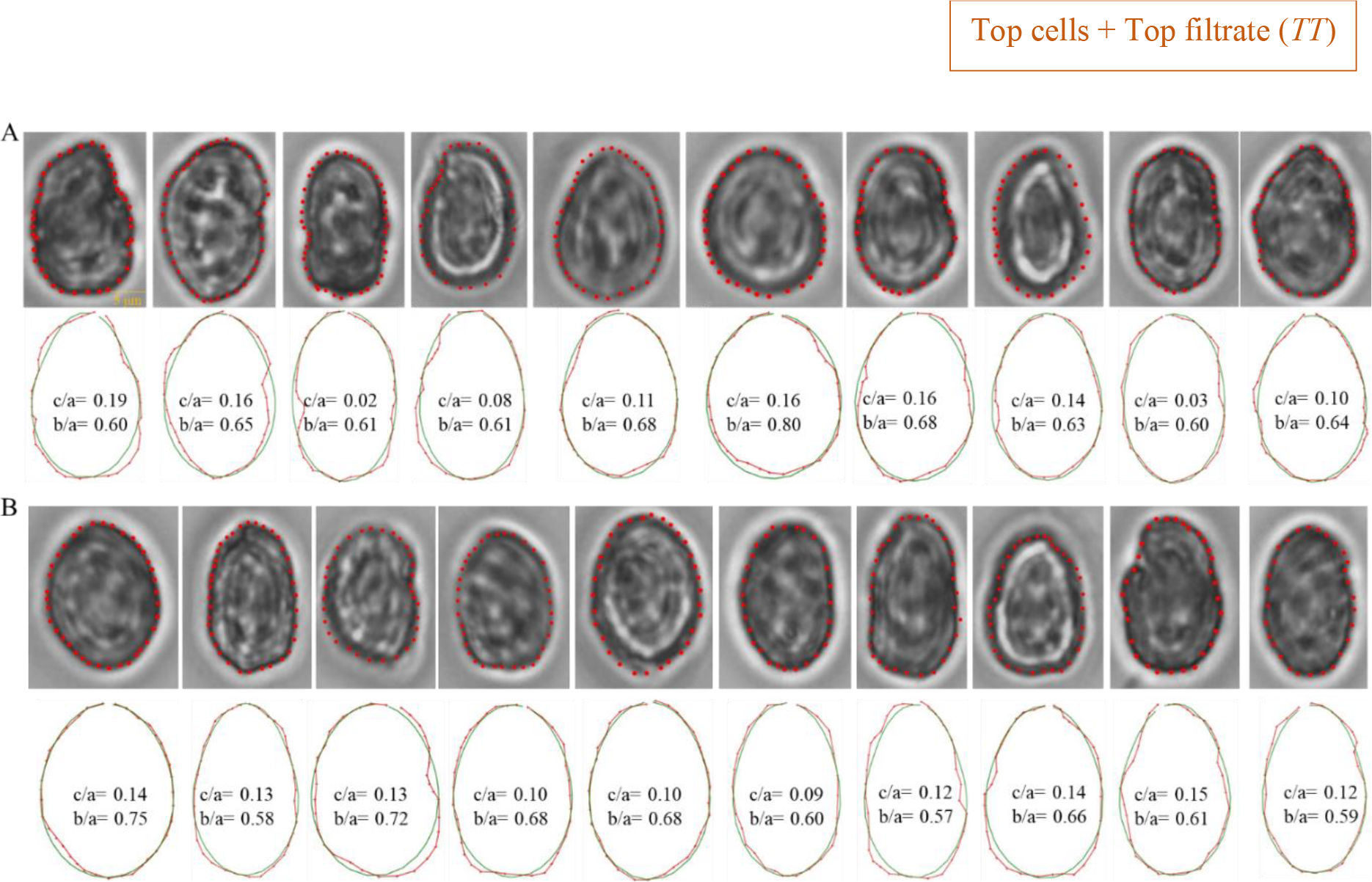
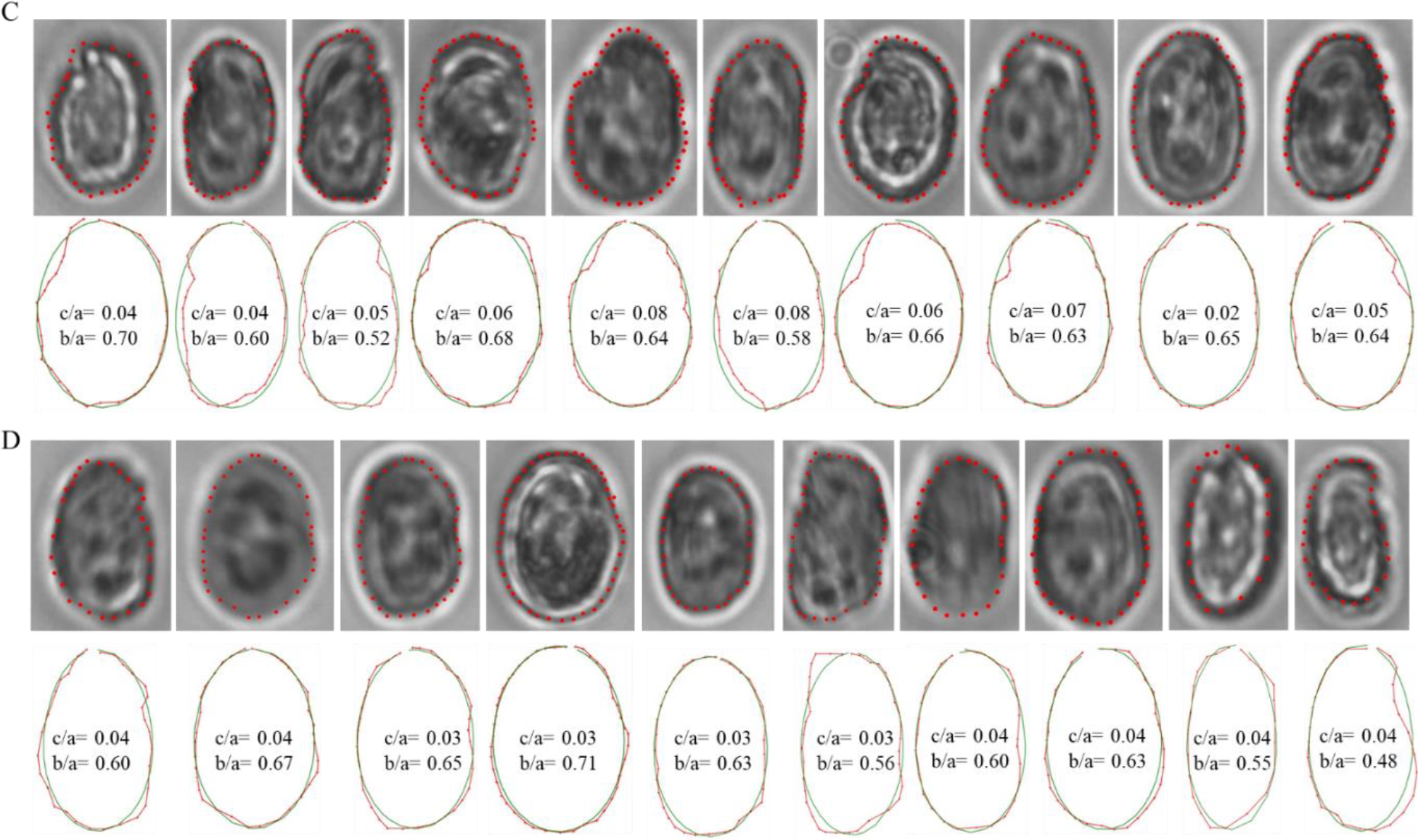
Quantification of morphological parameters of top cells in top filtrate [*T, T*]. The top row of each panel shows phase contrast images of top cells in presence of top filtrate (*N* = 10). The external contour of each cell was marked, and co-ordinates were extracted. These were then fed into an image analysis pipeline to fit each contour with a fore-aft asymmetric ellipse. The *c/a* parameter is a quantification of this asymmetry. Panels A to D represents distinct time points of 0, 20, 60 and 120 min.

**Fig. S12.**
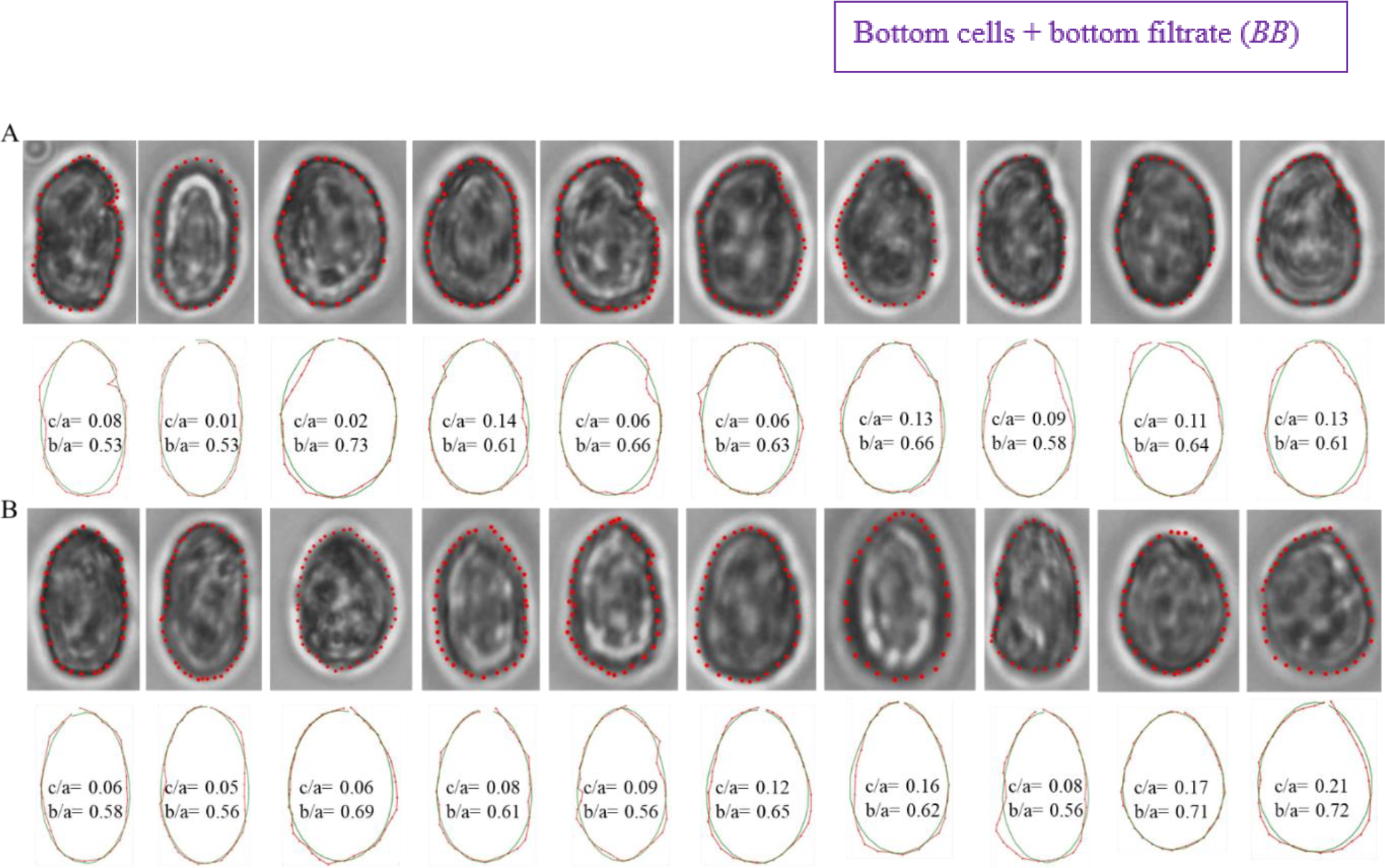
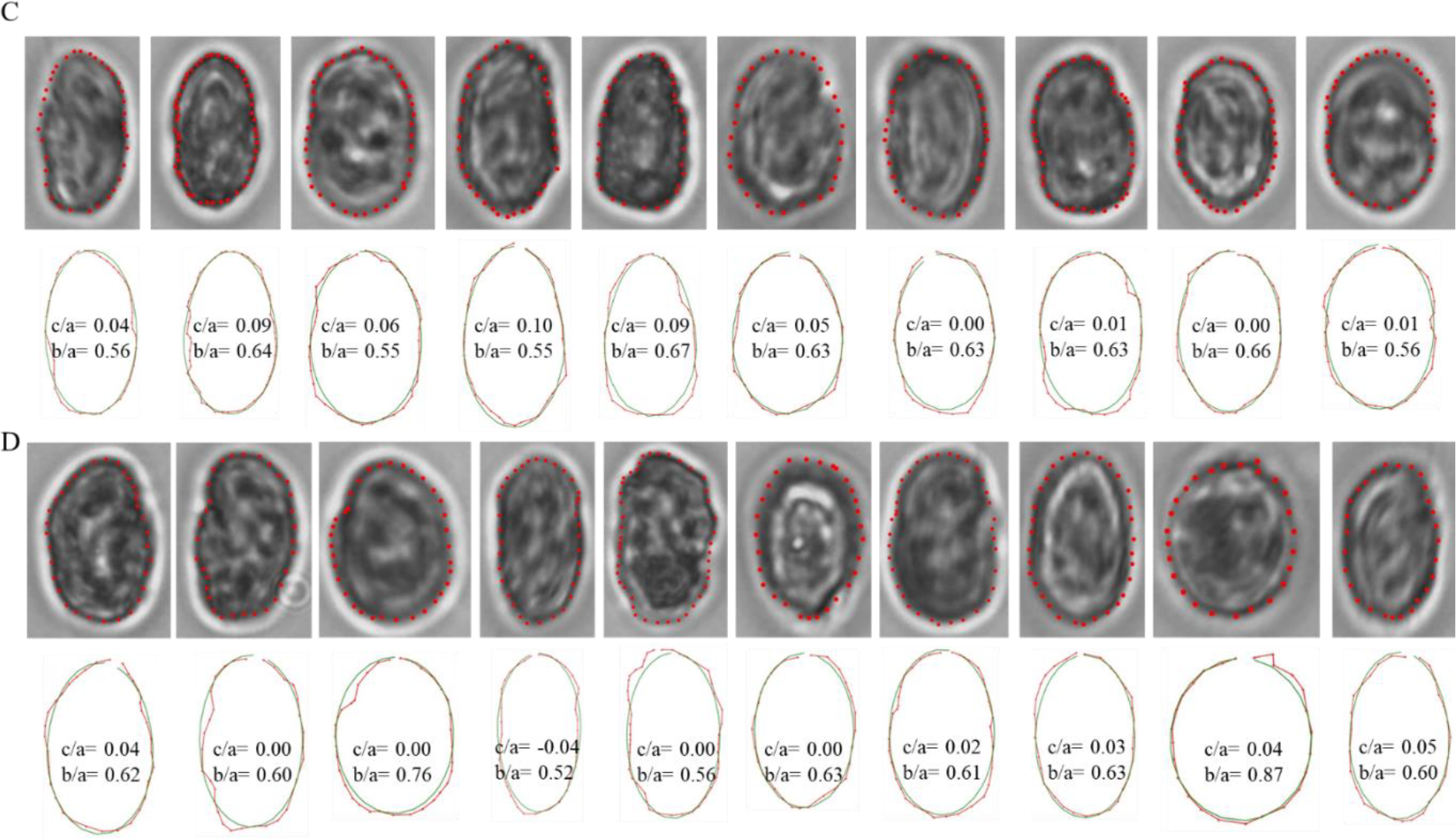
Quantification of morphological parameters of bottom cells in bottom filtrate *[B, B]*. The top row of each panel shows phase contrast images of top cells in presence of top filtrate (*N* = 10). The external contour of each cell was marked, and co-ordinates were extracted. These were then fed into an image analysis pipeline to fit each contour with a fore-aft asymmetric ellipse. The *c/a* parameter is a quantification of this asymmetry. Panels A to D represents distinct time points of 0, 20, 60 and 120 min.

**Fig. S13.**
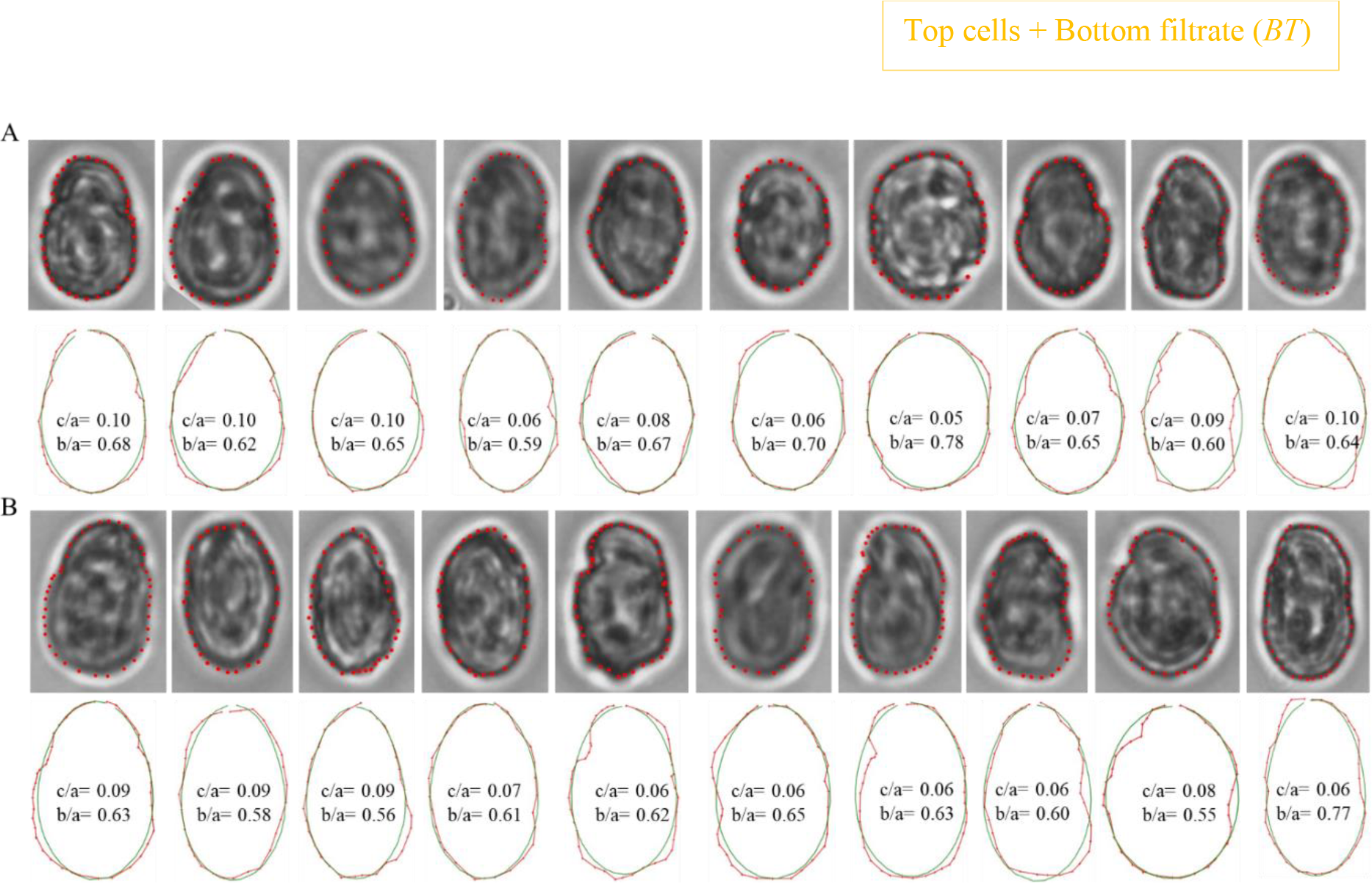
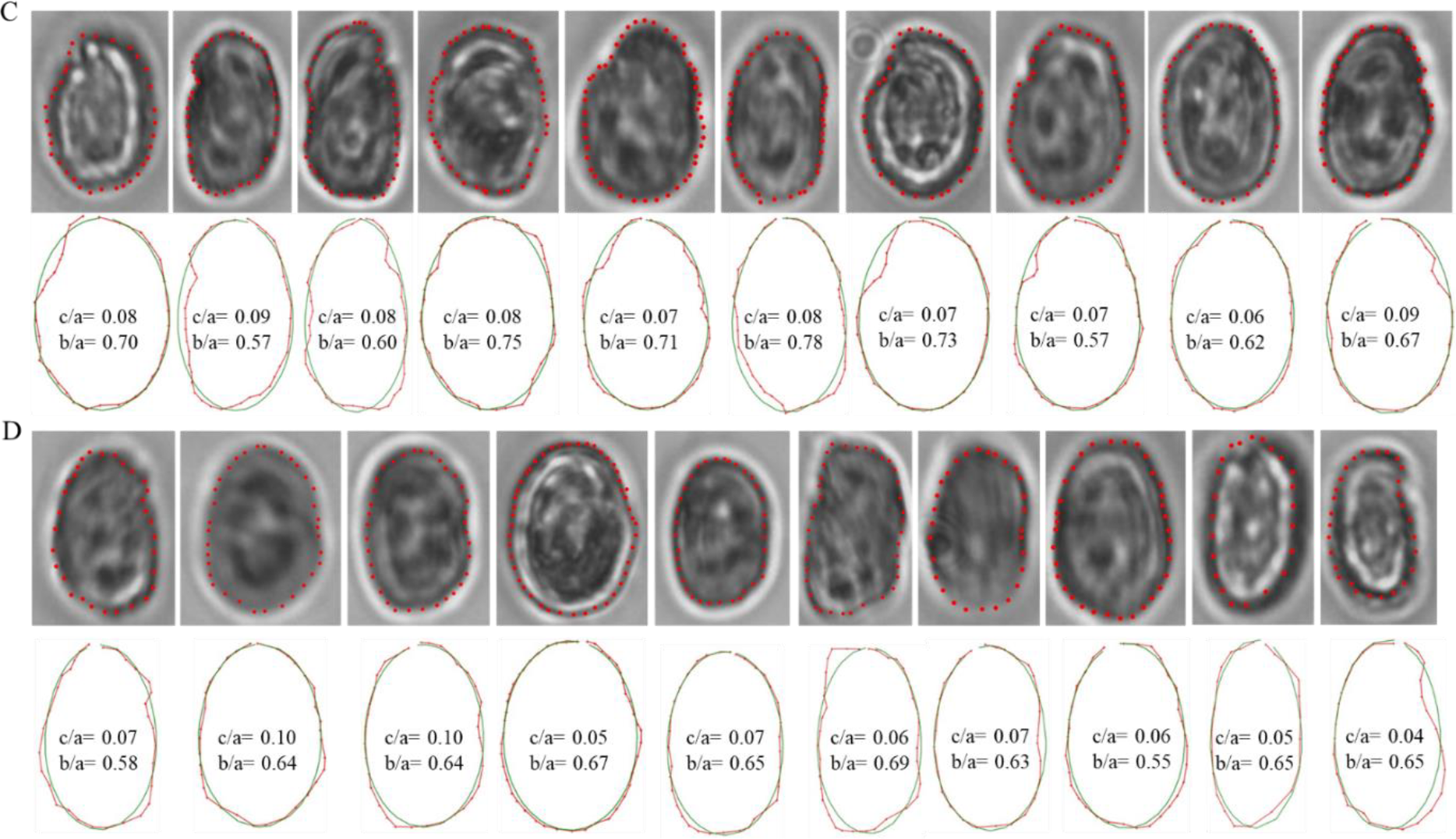
Quantification of morphological parameters of top cells in bottom filtrate *[B, T]*. The top row of each panel shows phase contrast images of top cells in presence of top filtrate (*N* = 10). The external contour of each cell was marked, and co-ordinates were extracted. These were then fed into an image analysis pipeline to fit each contour with a fore-aft asymmetric ellipse. The *c/a* parameter is a quantification of this asymmetry. Panels A to D represents distinct time points of 0, 20, 60 and 120 min.

**Fig. S14.**
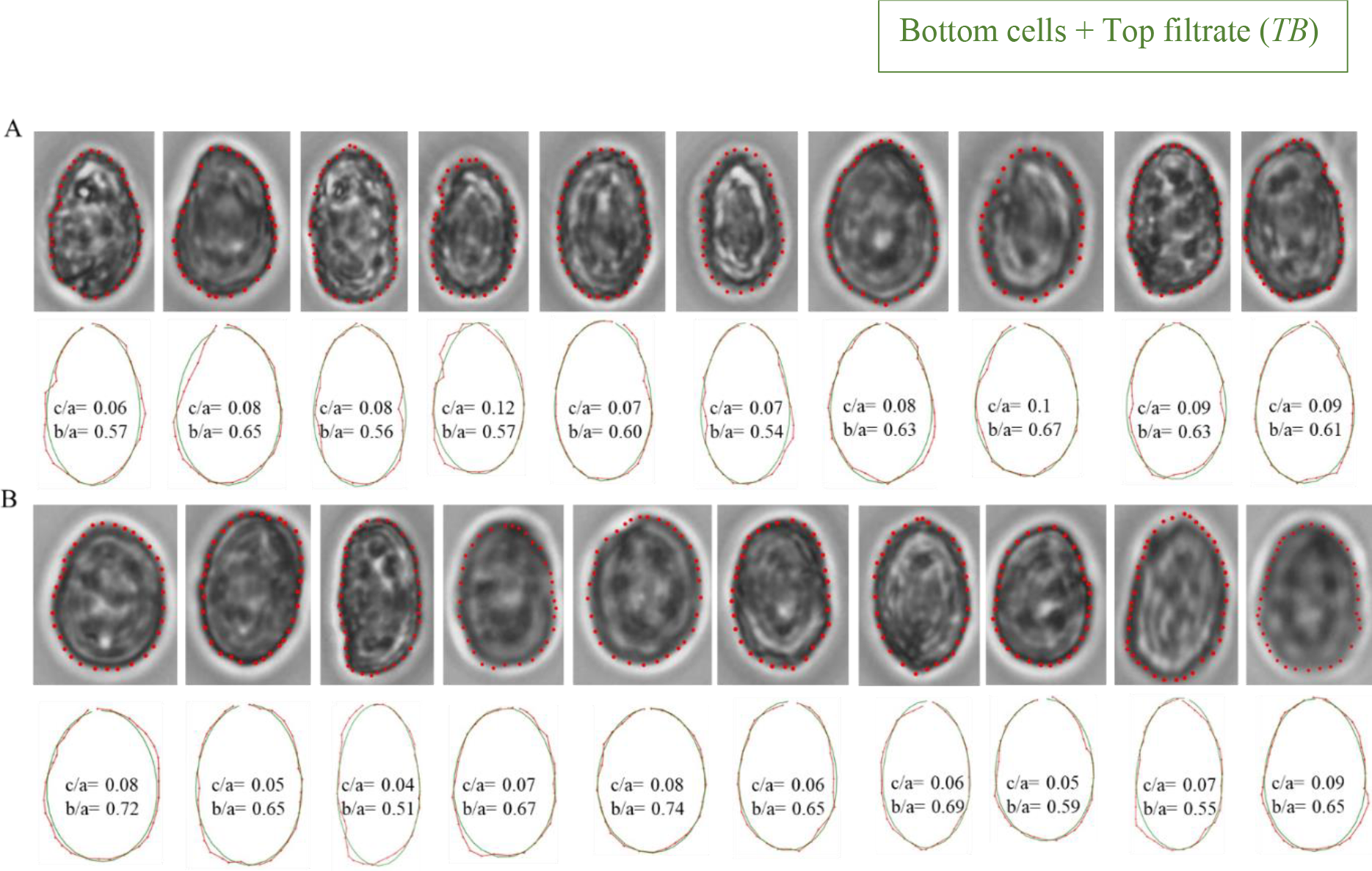
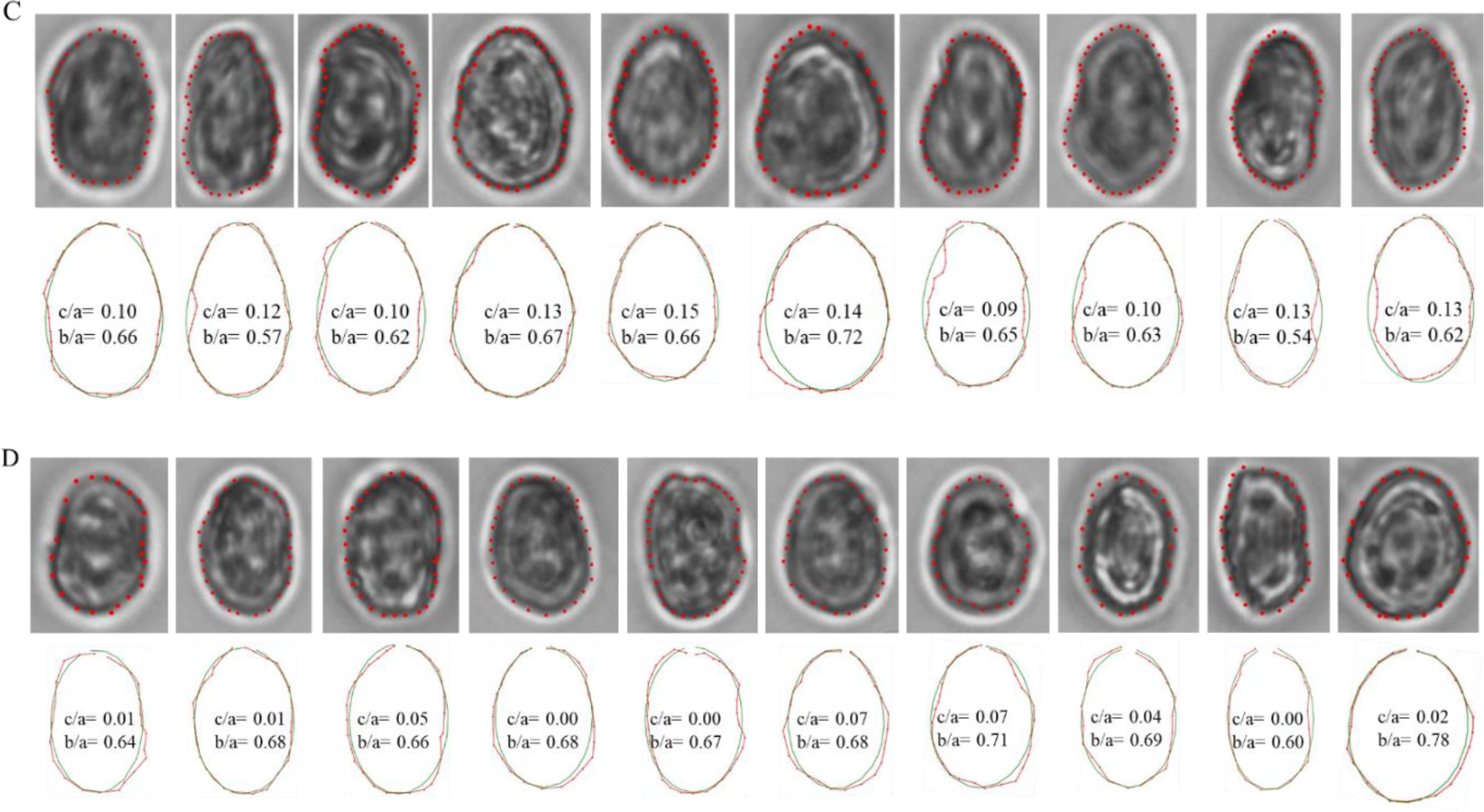
Quantification of morphological parameters of bottom cells in top filtrate *[T, B]*. The top row of each panel shows phase contrast images of top cells in presence of top filtrate (*N* = 10). The external contour of each cell was marked, and co-ordinates were extracted. These were then fed into an image analysis pipeline to fit each contour with a fore-aft asymmetric ellipse. The *c/a* parameter is a quantification of this asymmetry. Panels A to D represents distinct time points of 0, 20, 60 and 120 min.

**Fig. S15.**
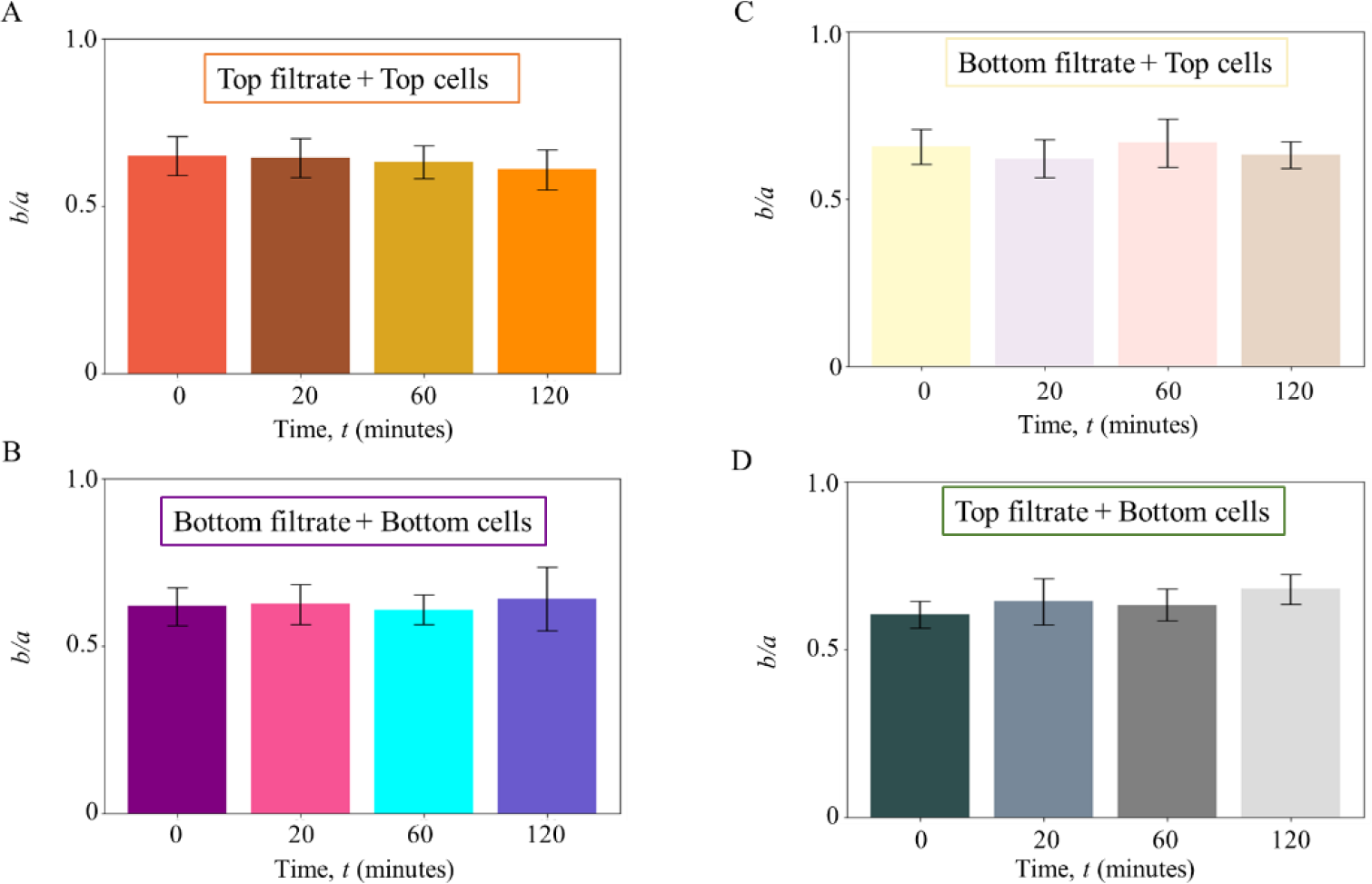
*b/a* values of cells over time in presence of filtrates. **(A)** For top cells in presence of top filtrate *[T, T]*, the mean (± s.d.) values of *b/a* over 0, 20, 60 and 120 minutes are 0.65 ± 0.06, 0.64 ± 0.06, 0.63 ± 0.05 and 0.61 ± 0.06. **(B)** For bottom cells in presence of bottom filtrate *[B, B]*, the mean (± s.d.) values are 0.62 ± 0.06, 0.62 ± 0.06, 0.61 ± 0.05 and 0.64 ± 0.1. **(C)** For top cells in presence of bottom filtrate *[B, T]*, the mean (± s.d.) values are 0.66 ± 0.05, 0.62 ± 0.06, 0.67 ± 0.07 and 0.63 ± 0.04. **(D)** For bottom cells in presence of top filtrate *[T, B]*, the mean (± s.d.) values are 0.6 ± 0.04, 0.64 ± 0.07, 0.63 ± 0.05 and 0.68 ± 0.04.

**Fig. S16.**
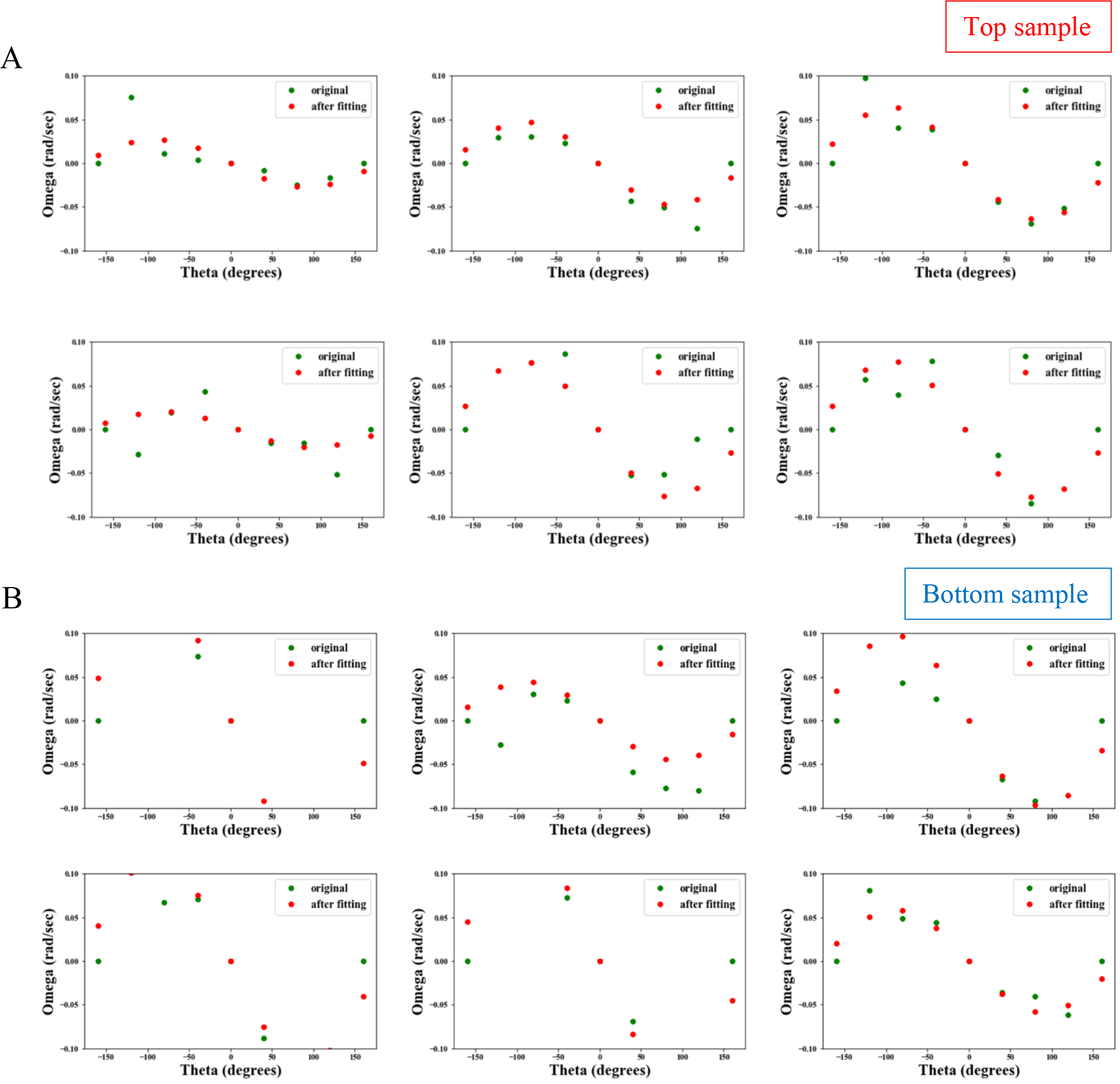
Quantification of the orientational stability of cell samples during night. To compute the stability of top and bottom samples, rotation rate (ω) of cells were plotted as a function of the instantaneous angular direction (*θ*). The time of experiment was 00:00 h. **(A)** shows the stability of top cells with each panel corresponding to one replicate (of 6 replicates, as mentioned in Fig. S3). Red dots correspond to experimental data and green dots to the sinusoidal curve obtained by fitting a sinusoid to the experimental data. **(B)** shows the same but for bottom cells. The amplitude, *A* of the fitted curve was used to obtain the reorientation timescale, *B* as 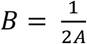.

**Fig. S17.**
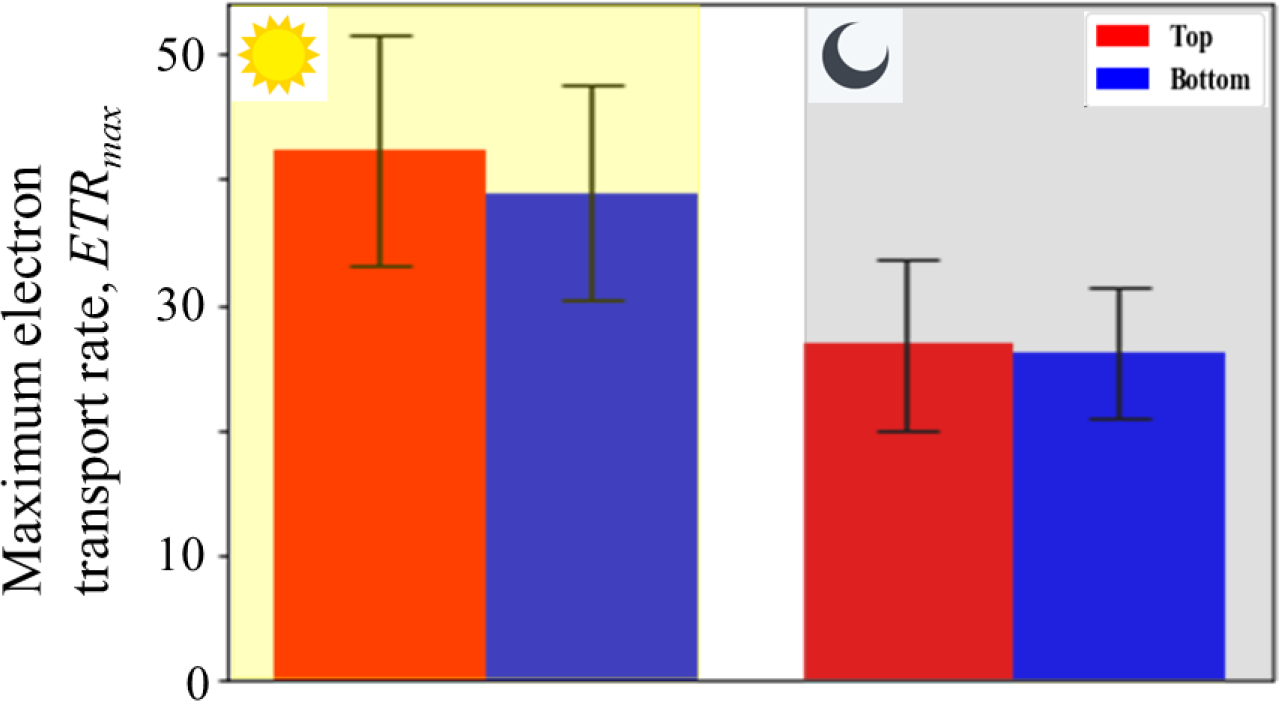
Estimation of the electron transport rate (*ETR*_*max*_) using the PAM device. Complementing the maximum photosynthetic yield and non-photochemical quenching, two key processes during light harvesting, this figure shows the maximum ETR, *ETR*_*max*_, during the same. During daytime, the difference between top and bottom for this value is not statistically significant (p = 0.69). At night as well, there is no significant difference (p = 0.94).

**Fig. S18.**
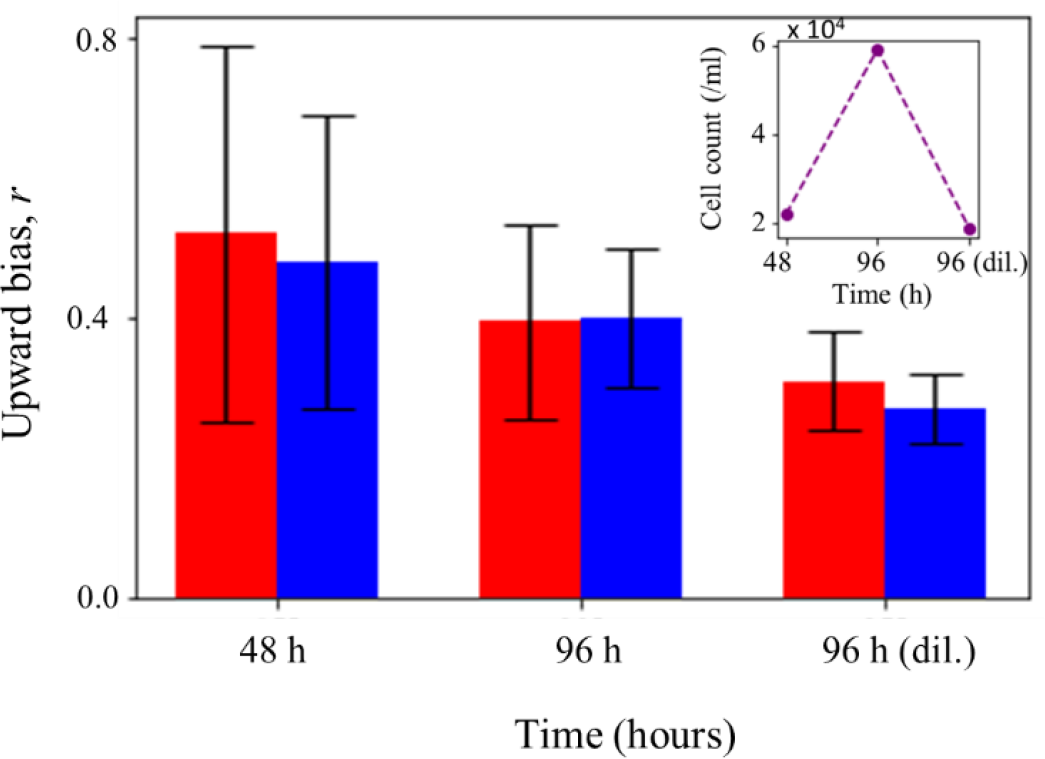
Effect of cell concentration on swimming behavior. To study the effect of concentration on population behavior, two points in growth phase: 48 h and 96 h were chosen for which the upward bias (*r*) values were measured (cell counts are shown in the inset). At 48 h, both top (red bar) and bottom (blue bar) have higher mean (± s.d.) *r* than at 96 h. Same cells from 96-hour culture were taken and diluted (dil.) ∼3 times with corresponding filtrate, to achieve a concentration comparable to that at 48 h (third data point, inset). The *r* value of this diluted sample was calculated and compared to the other two datasets. 4 replicates were used for 48 h and 96 h (diluted), and 6 at the 96 h timepoint.

## Notes

**Funding:** This work was supported by the Luxembourg National Research Fund’s AFR-Grant (Grant no. 13563560) and the ATTRACT Investigator Grant, A17/MS/11572821/MBRACE, to A.S. Support from the German Science Foundation project (DFG, Pycnotrap GR1540/37-1 to H-P.G.) is gratefully acknowledged.

### Competing Interest Statement

The authors have declared no competing interest.

### Summary of Updates

1) New figures, data and table in Supplementary section 2) Discussions and conclusions udpated 3) Affiliation of author-2 updated 4) Correction of typographical errors

